# The BrightEyes-TTM: an Open-Source Time-Tagging Module for Single-Photon Microscopy

**DOI:** 10.1101/2021.10.11.463950

**Authors:** Alessandro Rossetta, Eli Slenders, Mattia Donato, Eleonora Perego, Francesco Diotalevi, Luca Lanzanó, Sami Koho, Giorgio Tortarolo, Marco Crepaldi, Giuseppe Vicidomini

## Abstract

Fluorescence laser-scanning microscopy (LSM) is experiencing a revolution thanks to the introduction of new asynchronous read-out single-photon (SP) array detectors. These detectors give access to an entirely new set of single-photon information typically lost in conventional fluorescence LSM, thus triggering a new imaging/spectroscopy paradigm – the so-called single-photon LSM (SP-LSM). The revolution’s outcomes are, from one side, the blooming of new SP-LSM techniques and tailored SP array detectors; from the other side, the need for data-acquisition (DAQ) systems effectively supporting such innovations. In particular, there is a growing need for DAQ systems capable of handling the high throughput and high temporal resolution information generated by the single-photon detectors. To fill this gap, we developed an open-source multi-channel time-tagging module (TTM) based on a field-programmable-gate-array (FPGA), that can temporally tag single-photon events – with 30 ps precision – and synchronisation events – with 4 ns precision. Furthermore, being an open-access project, the TTM can be upgraded, modified, and customized by the microscopy-makers. We connected the TTM to a fluorescence LSM equipped with a single-photon avalanche diode (SPAD) bi-dimensional array detector, and we implemented fluorescence lifetime image scanning microscopy (FLISM) and, for the first time, fluorescence lifetime fluctuation spectroscopy (FLFS). We expect that our BrigthEyes-TTM will support the microscopy community to spread SP-LSM in many life science labs.

## Main

A revolution is happening in fluorescence laser scanning microscopy (LSM): the radically new way in which the fluorescence signal is recorded with single-photon array detectors drastically expands the information content of any LSM experiment.

For a conventional fluorescence laser scanning microscope, an objective lens focuses the laser beam at a specific position in the sample, the so-called excitation/probing region. The same objective lens collects the emitted fluorescence and, together with a tube lens, projects the light onto the sensitive area of a single-element detector, such as a photo-multiplier tube (PMT). The probing region is scanned across the sample or kept steady depending on the type of experiment: For example, for imaging is raster scanned, for single-point fluorescence correlation spectroscopy (FCS) is kept steady, for scanning FCS (sFCS) is scanned repeatedly in a circular or linear manner. During the whole measurement, the detector generates a one-dimensional signal (intensity versus time), that the data-acquisition (DAQ) system integrates within the pixel-dwell time (for imaging) or within the temporal bins (for FCS). Such a signal recording process induces information loss: the fluorescence light is integrated regardless of its position on the sensitive area and its arrival time with respect to a particular event, such as the excitation event. Notably, also other properties of light, such as the wavelength and the polarisation, are typically completely or partially discarded.

New single-photon (SP) array detectors, when combined with adequate DAQ systems, have the possibility to preserve most of this information. In particular, asynchronous read-out SP array detectors (a matrix of fully independent elements able to detect single photons with several tens of picoseconds timing precision) have made it possible to spatiotemporally tag each fluorescence photon, i.e., to record simultaneously at which position of the array (spatial tag) and at which delay with respect to a reference time (temporal tag) the photon hits the detector.

At present, the spatial tags can be used in two ways: firstly, they allow imaging of the probing region by placing a bi-dimensional detector array in the LSM image plane. Secondly, by dispersing the fluorescence across the long axis of a linear detector array, the spatial tags enable spectrally-resolved recording of the probing region, i.e., the spatial tag encodes the wavelength of the photon. At the same time, by exciting the sample with a pulsed laser and recording the fluorescence photon arrival time with respect to the excitation event, the temporal tags (i.e., the time difference between the excitation event and the photon detection) allow for sub-nanosecond time-resolved measurements, such as fluorescence lifetime or photon correlation. Furthermore, recording the photon arrival times with respect to the beginning of the experiment allows for microsecond intensity fluctuation analysis (i.e, microsecond time-resolved spectroscopy). In summary, these spatial, temporal, and spectral photon signatures have opened a series of advanced fluorescence imaging and spectroscopy techniques precluded to (or made more complex by) conventional single-element detectors. Hereafter, we refer to these new techniques with the collective term single-photon laser-scanning microscopy (SP-LSM).

Very recently, a new LSM architecture based on a SP linear detector has led to a revival of the combination of fluorescence lifetime and spectral imaging (1) – spectral fluorescence lifetime imaging microscopy (S-FLIM). Simultaneously, bi-dimensional SP array detectors have opened up new perspectives for image-scanning microscopy (ISM). In a nutshell, ISM uses the information contained in the spatial distribution (image) of the probing region to reconstruct the specimen image with a twofold spatial resolution and a higher signal-to-noise ratio (SNR) than conventional LSM (2–4). Because SP bi-dimensional array detectors provide a sub-nanosecond time-resolved image of the probing region, ISM can be combined with fluorescence lifetime to create a super-resolution fluorescence lifetime imaging technique, called fluorescence lifetime ISM (FLISM) (5), or to trigger the implementation of a new class of nanoscopy techniques, namely quantum ISM (Q-ISM) (6). The microsecond time-resolved images can also be used to implement (i) high information content FCS and, more generally, fluorescence fluctuation spectroscopy (FFS) (7) – usually referred to as comprehensive-correlation analysis (CCA) (8); and (ii) the combination of super-resolution optical fluctuation imaging (SOFI) with ISM (9) (SOFISM).

Key elements for implementing SP-LSM are the SP array detector and the DAQ system. Although analogue-to-digital converters (e.g., constant-fraction discriminators) allow the use of photo-multiplier tube arrays as SP detectors, they introduce a significant amount of unwanted correlation into the measurements (10) and are outperformed by true SP detectors for what concerns the photon-time jitter/precision. An alternative to PMT-based SP array detectors is the AiryScan-like module (11), a hexagonal-shaped multi-cores fiber bundle connected to a series of single-element avalanche-photon diodes (APDs) (6). Clearly, this module is highly expensive and not scalable. True single-photon array detectors based on the well-established SPAD array technology (12, 13) solved these limitations. In particular, new asynchronous read-out SPAD array detectors have been tailored for SP-LSM applications (14–16), i.e., these detectors have small number of elements – but enough for sub-Nyquist sampling of the probing region – high photon-detection efficiency, low dark-noise, high dynamics, high fill-factor, low cross-talk, and low photon time-jitters. Furthermore, SPAD array detectors offer, in general, a low cost, a high robustness and microsized dimensions. Whilst the development of SPAD array detectors specifically designed for SP-LSM is gaining substantial momentum, no effort has been placed in developing an open-source data acquisition (DAQ) systems able to (i) fully exploit the high-throughput and high-resolution photon-level information that these detectors provide; (ii) offer flexibility and upgradability. The lack of an open-source DAQ system may preclude a massive spreading of the above SP-LSM techniques and the potential a blooming of new effective imaging and spectroscopy techniques based on this new SP-LSM paradigm.

In this manuscript, to address this need, we propose an open-source multi-channel time-tagging module (TTM), the BrightEyes-TTM, specifically designed to implement current and future SP-LSM techniques. In short, the BrightEyes-TTM implements multiple photon- and reference-channels to record at which element of the detector array, and when (with tunable precision), in respect to the reference/synchronisation events, the single-photon reaches the detector. The BrightEyes-TTM is based on a commercially available and low cost field-programmable-gate-array (FPGA) evaluation board, equipped with a state-of-the-art FPGA and a series of I/Os connectors granting an easy interface of the board with the microscope, the SPAD array detector and the computer. We chose the FPGA to grant both quick prototyping and versatility, and to meet the future requests from new SP-LSM applications/techniques.

Our implementation was conceived and designed having in mind the fluorescence applications and the SPAD array detector characteristics, current and future, thus providing a good compromise between cost, photon-timing precision and resolution, temporal range, electronics dead-time, maximum photon-flux. Additionally, the low FPGA resource needed by the current implementation ensures the highly scalability of the architecture, potentially enabling more channels and, in general, new applications.

We integrated the BrightEyes-TTM into an existing SP-LSM equipped with a 21-element SPAD array detector and we used it to perform FLISM imaging on a series of calibration and biological samples. Furthermore, for the first time, we combined CCA with fluorescence lifetime, thus triggering a new series of fluorescence lifetime fluctuation spectroscopy (FLFS) techniques. Despite the great potential of SP-LSM, we are aware that a massive dissemination of this imaging paradigm will be effective only if a broad range of laboratories will have access to the TTM, and potentially modify it according to their needs. For this reason, this manuscript provides detailed guidelines, hardware parts lists, and opensource code for the FPGA firmware, and operational software.

## Results

### Multi-Channel Time-Tagging Module

To characterise the performances of the BrightEyes-TTM, we first used a test bench architecture based on the SYLAP pulse generator. In particular, we used the test bench to validate the fine (picoseconds precision) time-to-digital converters (TDCs) in measuring the start-stop time. Independently, we validated the coarse (nanosecond precison) TDCs directly by integrating the TTM into a SP-LSM system.

We measured the linearity of the fine TDC by performing a statistical code-density test: we fed a fixed-frequency (50 MHz) signal into the synchronisation (SYNC) channel and a random signal in one of the photon channels (here, channel 11). After accumulating several millions of photon events, we built the start-stop time histogram, also called the time-correlated-single-photon counting (TCSPC) histogram, which shows in a differential non-linearity (DNL) of *σ*_DNL_ = 0.06 least-significant-bit (LSB) and an integral non-linearity (INL) of *σ*_INL_ = 0.08 LSB (Fig. 1 a) with LSB = 48 ps.

**Fig. 1.**
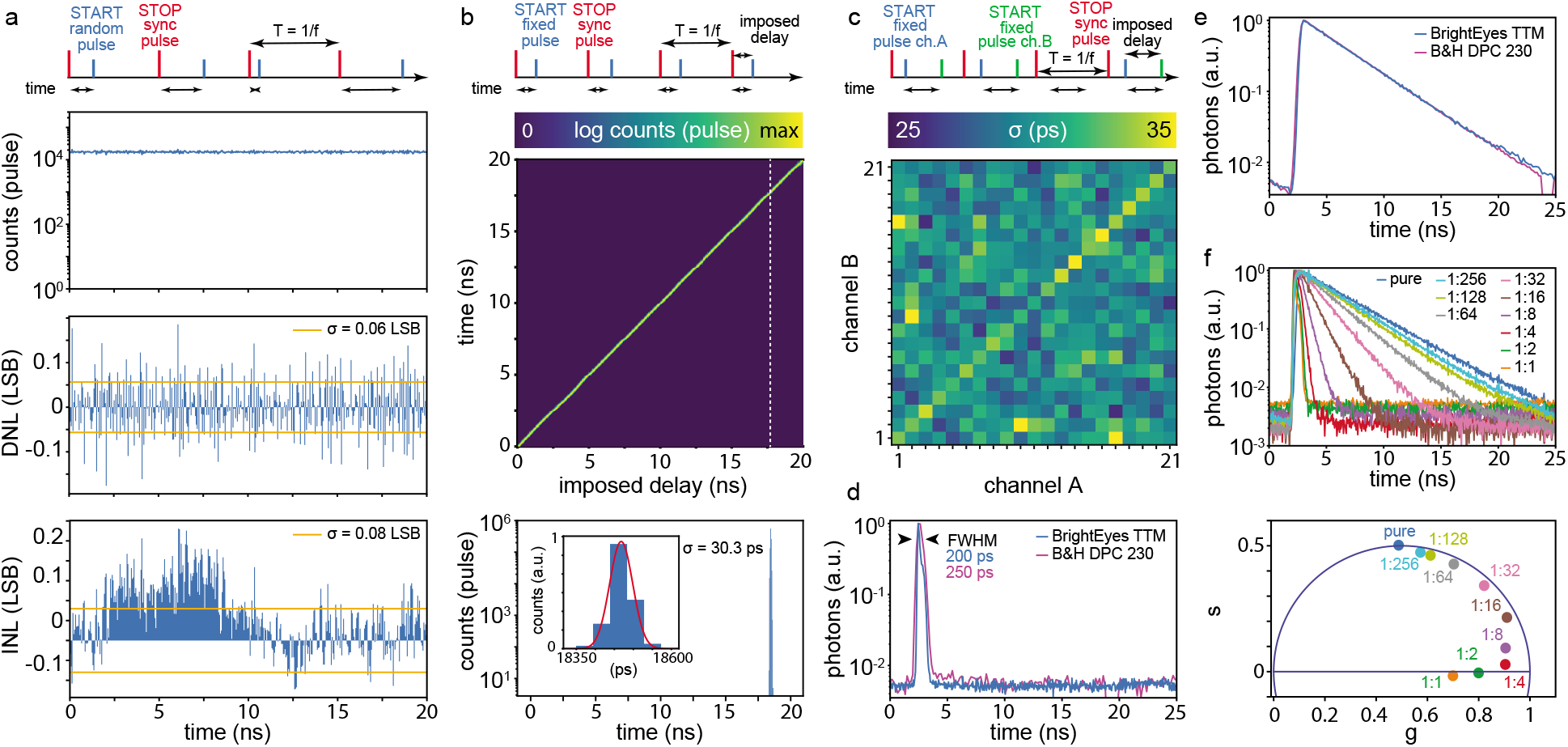
BrightEyes-TTM characterisation and validation. **a** Statistical code-density test. Temporal schematic representation for the signals involved in the experiment: a fixed frequency SYNC (STOP) clock signal at *f* = 50 MHz (i.e, *T* = 1/*f* = 20 ns) and an uncorrelated/random train of pulses (START). The reconstructed start-stop time histogram, i.e., counts versus time (top), the relative DNL (middle) and INL (lower). **b** Single-shot precision experiment. Temporal schematic representation: a fixed frequency SYNC clock signal and a synchronised but delayed (in a controlled way) signal. Unified representation of the start-stop time histograms as a function of the imposed delay between the two signals (top). Single start-stop time histogram for the delay denoted by the dotted white line in the middle pane (bottom). The inset shows a magnification of the histogram for a selected temporal interval, superimposed with the Gaussian fit (red line). **c** Dual-channel single-shot precision experiment. Temporal schematic representation: a fixed frequency SYNC clock signal and a pair of synchronised signals (channel A and channel B). The delays between all three signals are fixed. Jitter map for each pair of channels, i.e., error in the time-difference estimation between any two channels, measured as the standard deviation of a Gaussian fit of the error distribution. The diagonal of the map represents the sigma of the single-channel single-shot precision experiment. **d** Normalised impulse response functions (and FWHM values) for the BrightEyes-TTM and DPC-230 multi-channel card. The IRFs represent the response of the whole architecture (microscope and DAQ) to a fast (sub-nanosecond) fluorescent emission. **e** Normalised fluorescence decay histogram (i.e., start-stop time histogram or TCSPC histogram) for a fluorescein-water solution measured with the BrightEyes TTM and the DPC-230 card. **f** Normalised fluorescence lifetime decay histograms of quenched fluorescein solutions for increasing concentrations of quencher (potassium iodide) (top), phasor representation of quenched fluorescein solutions (bottom). All single-channel measurements were done with TTM channel #10, which received the photon signal from the central element of the SPAD array detector. All start-stop (i.e., TCSPC) histograms have 48 ps resolution (bin-with).

We then measured the single-shot precision (SSP) of the fine TDC by measuring repeatedly a constant start-stop time interval: we fed a fixed-frequency (50 MHz) signal into the SYNC channel and a synchronised second signal – with a tunable fixed delay – into one of the photon channels (here, channel 11). After accumulating several millions of sync-photon pairs, we built the start-stop time histogram, which in this case represent the distribution of the measurement error. (Fig. 1 b). By fitting the start-stop time histogram with a Gaussian function, we estimated a precision of *σ* = 30 ps. We tuned the delay between the two signals across the whole temporal of the fine TDC (here, 20 ns) and we always observed a similar precision (Suppl. Fig. S1), confirming the excellent linearity of the fine TDC. We repeated the same SPP experiment for all the photon channels and we obtained a similar precision (diagonal values on matrix of Fig. 1 c).

Furthermore, to confirm the ability of the 21 channels to record photons in parallel, we repeated the SSP experiment by feeding the same signal also into a second photon channel. In this case, the delays between all three signals (Channel_*A*_, Channel_*B*_, and SYNC) were kept fixed, and we used the TTM to measure the delay between the two photon channel signals. We built the histogram, similar to the start-stop time histogram, that reports the elapsed time between the two photon channel signals, and we fit the distribution with a Gaussian function. We performed the experiment for all the possible channel pairs and we obtained precision values of *σ* = 25÷35 ps, depending on the channels pair (Fig. 1 c).

Next, we integrated the BrightEyes-TTM into a custom-built single-photon laser scanning microscope equipped with a bi-dimensional 21-element SPAD array detector and a picosecond pulsed diode laser. We used a quenched solution of fluorescein in water, saturated with potassium iodide to measure the impulse-response function (IRF) of the system (17). The relatively large full-width-at-half-maximum value of the IRF (240 ps) is due to the convolution of the single-shot response (~ 30 ps) with the laser pulse-width (*>* 100 ps), the SPAD photon jitters (~ 120 ps) and the jitters/dispersion introduced by the optical system (Fig. 1 d). We compared the IRF of the BrightEyes-TTM with the IRF of the DPC-230 commercial multi-channel time-tagging card. Notably, because of the poor time resolution (164 ps from data-sheet), the DPC-230 is not able to reveal the typical cross-talk effect of the SPAD array detector (16), which is visible in the BrightEyes-TTM as an additional bump (Fig. 1 d). We used the two time-tagging systems to compare the decay distributions of a pure (not quenched) solution of fluorescein in water. The two TCSPC histograms show very similar shapes, which is confirmed by fitting them with a single exponential decay model (*τ_fl_* = 3.97 ± 0.04 ns and *τ_fl_* = 3.99 ± 0.01 ns, for the BrightEyes TTM and the DPC-230, respectively) (Fig. 1 e). To demonstrate the ability of the BrightEyes-TTM to work at different temporal s, we repeated the pure fluorescein experiment for different laser frequencies (80,40, 20, 10 and 5 MHz). The TCSPC histograms do not show significant differences (Suppl. Fig. S2).

To test the BrightEyes-TTM for different fluorescence lifetime values, we measured the decay distributions of the quenched fluorescein solution for increasing concentrations of potassium iodide (Fig. 1 f). The higher the potassium iodide concentration is, the higher the quenching will be, and the longer the measurement time needed to accumulate good photon statistics. For this reason, the dark-noise (which appear as an uncorrelated background in the TCSPC histogram) increases with the quencher concentration. The same effect appears on the phasor-plot: because the decays follow a single-exponential function, caused by the collisional mechanism of the quenching, all points, regardless of the concentration, should lie on the universal semicircle. However, the uncorrelated background shifts the points towards the origin, namely lower signal-to-background ratio yields an higher demodulation.

### Fluorescence Lifetime Image Scanning Microscopy

To demonstrate the ability of the BrightEyes-TTM in the context of SP-LSM imaging, we implemented FLISM on the same SP laser scanning microscope used for the IRF and fluorescein measurements. As with all LSM imaging techniques, FLISM requires recording the fluorescence photons in synchronisation with the scanning system (e.g., galvanometer mirrors, acoustic-optics deflectors, and/or piezo stages). We can obtain this synchronisation by measuring the photon arrival times with respect to the clock signals typically provided by the scanning system (i.e., pixel/line/frame clocks). Since the scanning synchronisation does not need high precision, we can use the reference (REF) channels of the TTM, which use a coarse TDC, i.e, nanosecond precision (~ 4.2 ns) to measure the relative delay between photon and clock signal. Notably, implementing a REF channel using a digital counter-based approach, thus a coarse TDC, consumes much less FPGA-resources than a high precision channel, which requires a fine TDC. Currently, the BrightEyes-TTM has three REF channels but could host more with minimal changes in the architecture. For example, more REF channels would allow to record the photons in synchronisation with a change in the excitation conditions, such as the intensity, the wavelength, or the polarisation. Indeed, the opto-mechanical devices used to change these experimental conditions (e.g., acousto optical modulators, shutters, and polarizer modulators) have a typical response time in the order of nanoseconds, compatible with the coarse TDC implementation.

We connected the pixel, line, and frame clocks delivered by the microscope controlling system to our TTM and we performed 2D imaging of phantom and biological samples. Thanks to the synchronisation signals (SYNC, pixel/line clocks), the stream of photons recorded by the TTM leads to a 4D photon-counting (intensity) image (*ch,x,y,τ*), where *ch* is the dimension describing the channel/element of the SPAD array, (*x, y*) are the spatial coordinates of the beam scanning system (in these experiments we recorded a single frame), and *τ* is the dimension of the TCSPC histogram (Table 3). First, we used the adaptive pixel-reassignment (APR) algorithm to reconstruct the 3D (*x,y,τ*) high spatial resolution and high signal-to-noise ratio ISM intensity image. Next, we applied both conventional fluorescence lifetime fitting and phasor analysis to obtain the FLISM (or lifetime) image (*x,y*) and the phasor plot/histogram (*g,s*), respectively. Side-by-side comparison of the intensity-based images clearly shows the optical resolution enhancement of ISM with respect to conventional (1.4 AU) imaging and the higher SNR with respect to confocal (0.2 AU) imaging (Fig. 2). The Fourier ring correlation (FRC) analysis confirms the effective resolution enhancement of ISM (Suppl. Fig. S3). In the context of fluorescence lifetime imaging, the higher SNR leads to an higher lifetime precision, as depicted by the lifetime histograms and the phasor plots (Fig. 2).

**Fig. 2.**
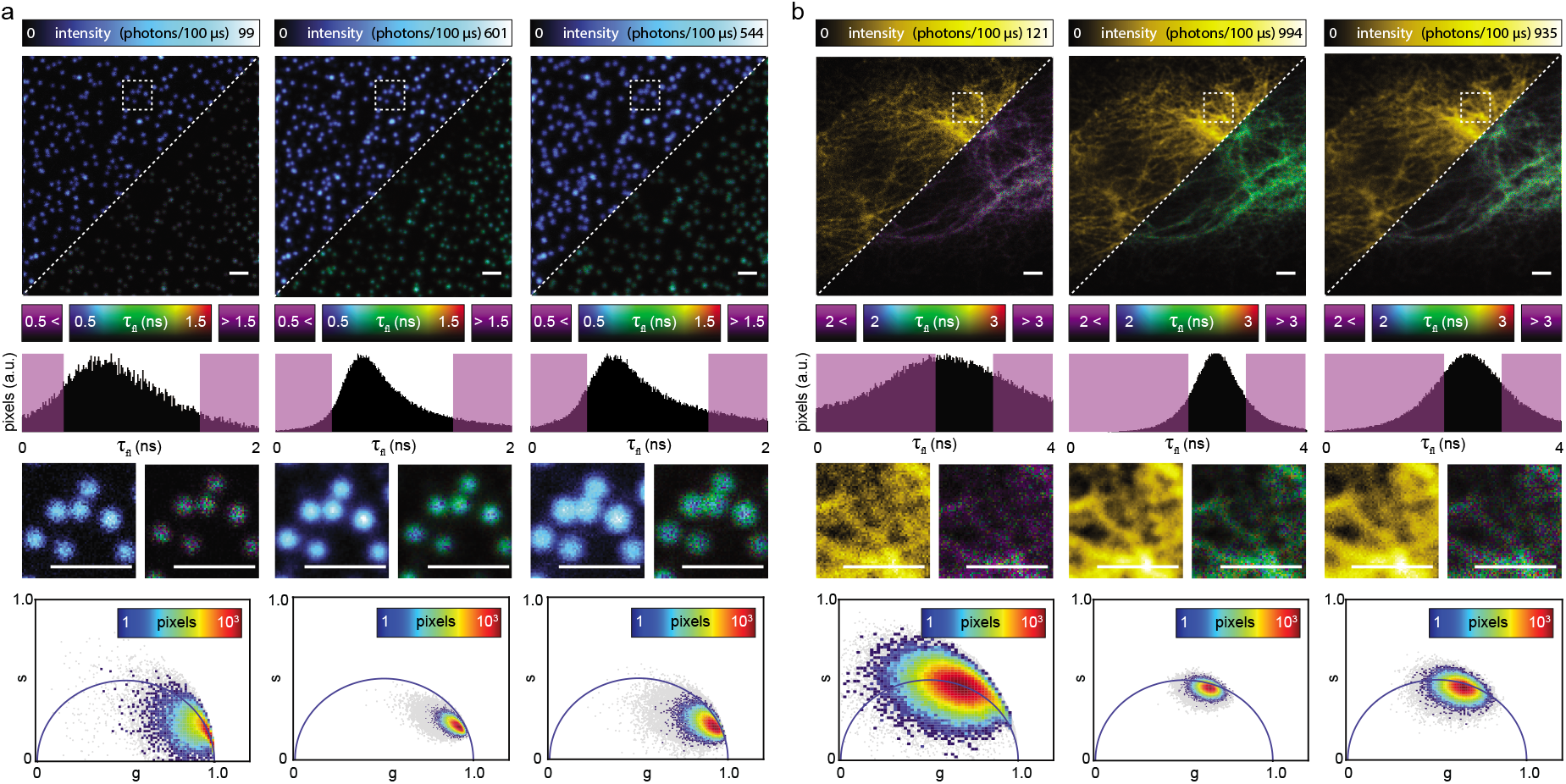
BrightEyes-TTM for FLISM. **a** Imaging and analysis of 100 nm nanobeads. Side-by-side comparison (top row) of closed confocal (left, pinhole 0.2 AU), adaptive pixel-reassignment ISM (center), and open confocal (right, pinhole 1.4). Each imaging modality shows both the intensity-based image (top-left corner) and the lifetime image (bottom-right corner). A bi-dimensional look-up-table is able to represent in the lifetime images both the intensity values (i.e., photon counts) and the excited-state lifetime values (i.e., *τ_fl_*). The intensity-based images integrate the relative 3D data (*x, y*, Δ*t*) along the start-stop time dimension Δ*t*. The lifetime-based images obtains the excited-state lifetime values by fitting the TCSPC histogram of each pixel with a single-exponential decay model. Histogram distributions of the imaging lifetimes values (middle-top row), number of pixels versus lifetime values, - in violet lifetime values which falls out of the selected lifetime representation interval. The lifetime images report in violet the pixels whose lifetime belong to this interval. Zoomed regions in the white-dash boxes, re-normalised to the maximum and minimum intensity values (middle-bottom). Pixel intensity thresholded phasor plots (5% and 10% thresholds respectively in gray and color). Phasor-plots were referenced using a fluorescein calibration sample. **b** Imaging and analysis of fluorescent labelled vimentin in fixed cell. Images and graphs appears in the same order as described for (a). Scale bars 2 *μ*m.

In order to obtain a real-time intensity-based image as a guideline during FLISM experiments, we implemented in the BrightEyes TTM an output digital signal which duplicates one of the photon channels (e.g, the channel of the central element of the SPAD array), and we sent it to the control system of the microscope (Suppl. Fig. S4). This approach was preferred to physically splitting the signal from one channel of the SPAD array, since the latter would likely alter the signal itself. Notably, this function represents an example of the versatility of the BrightEyes TTM in the context of imaging.

### Fluorescence Lifetime Fluctuation Spectroscopy

To show how the BrightEyes-TTM allows for the study of (bio)molecular dynamics, we implemented two types of spot-variation FLFS (18): circular scanning FLFS and steady-beam (i.e., single-point) FLFS. In both measurement types, the diffusion dynamics and the fluorescence lifetime can be obtained simultaneously from the absolute times (i.e., the delay of the photons in respect to the beginning of the experiment) *t*_photon_ and the start-stop times Δ*t*, respectively. Additionally, access to the start-stop times allows filtering the autocorrelation curves to mitigate artefacts such as detector afterpulsing.

As an example for the circular scanning FLFS, we measured freely diffusing fluorescent nanobeads (Fig. 3 (a-c)). From the absolute times, we calculated the unfiltered autocorrelations for the central SPAD array detector element, the sum of the 9 most centrals elements (called sum 3×3) and the sum of all the elements (called sum 5 × 5). By scanning the probing region in circles across the sample, the beam waist *ω*_0_ and the diffusion time *τ_D_* can be derived from the same experiment, i.e., no calibration measurement is needed. The fitted diffusion coefficient, (14.3 ± 0.5) *μm^2^/s*, corresponds to the value that can be expected based on the diameter of the beads and the diffusion law (or spot variation) analysis 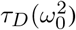 confirms that the beads were freely diffusing. However, the fitted concentration of beads does not scale proportionally to the focal volume, indicating that the amplitude of the autocorrelation functions are not correct. Using the start-stop time histograms, we calculated the filter functions which attenuate the detector afterpulsing and background signals. Applying the filters to the autocorrelations significantly alters the amplitudes, while the diffusion times are mostly left unchanged. As a result, the Δ*t*-based filtered autocorrelations show both the expected behaviour for the diffusion time and the concentration (Fig. 3 (a-c)). If the fluorescence signal is strong enough with respect to the background, filtering is not needed, as shown in (Fig. 3 (d)) for freely diffusing goat anti-mouse antibodies conjugated with Alexa 488. Here, the sample and the experimental conditions (e.g., dye concentration and laser power) lead to Δ*t* histograms without a significant uncorrelated background. As a result, the correlation curves do not need filtering, and both the diffusion time and the average number of particles in the focal volume follow the expected behaviour.

**Fig. 3.**
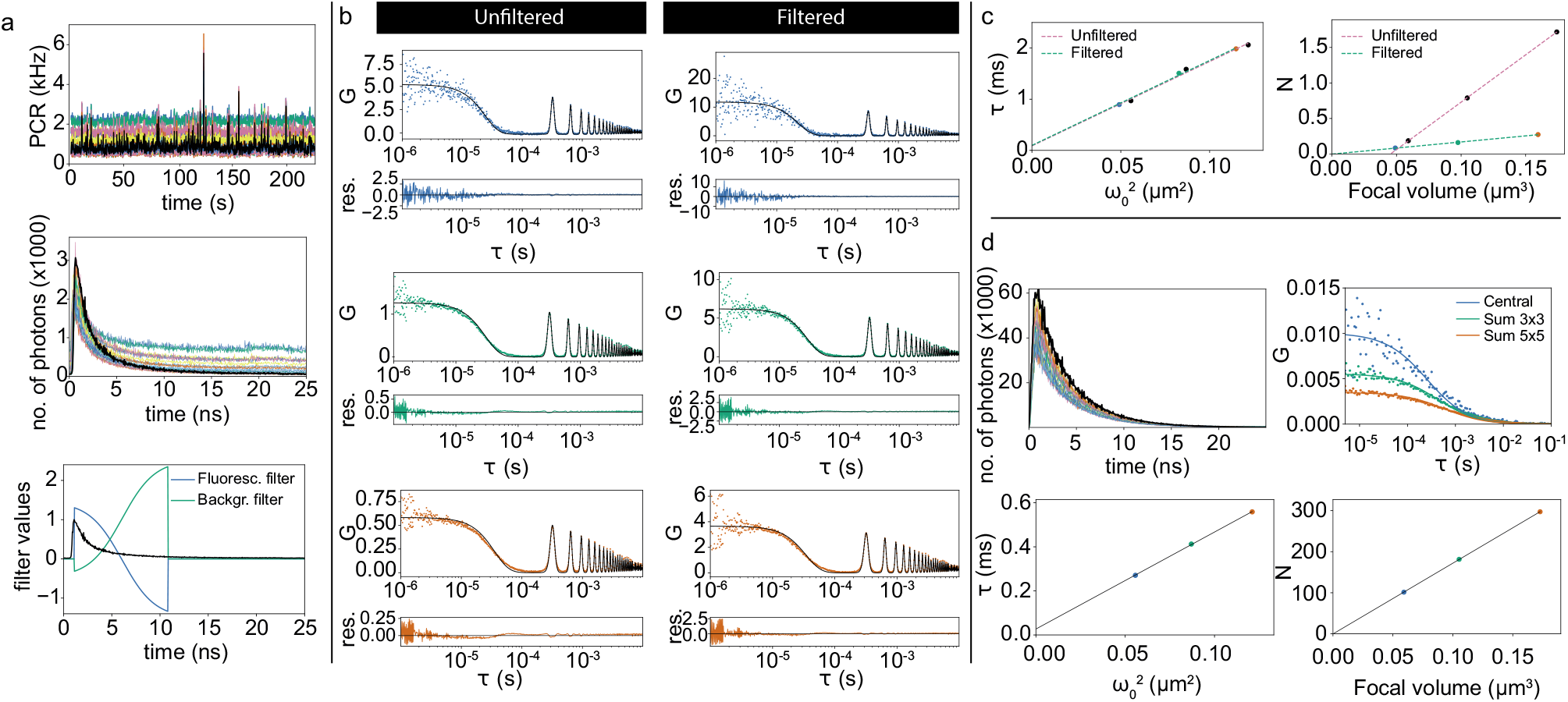
BrightEyes-TTM for FLFS on freely diffusing fluorescent beads (a-c) and goat anti-mouse antibodies conjugated with Alexa488 (d). **a** Photon count rate for all channels (top), start-stop time histograms (middle), and filter functions (bottom). The central pixel data is shown in black. **b** Autocorrelations and fits for the central pixel (top), sum 3×3 (middle), and sum 5×5 (bottom), for the unfiltered (left) and filtered (right) case. **c** Diffusion time as a function of 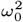 (left) and average number of particles in the focal volume as a function of the focal volume (right). The corresponding diffusion coefficients are (14.3 ± 0.5)*μm^2^/s* (unfiltered) and (14.0 ± 0.4)*μm^2^/s* (filtered). The fitted particle concentrations are (7 ± 3) /*μm*^3^ (unfiltered) and (1.70 ± 0.03) /*μm*^3^ **d** start-stop time histograms, (unfiltered) autocorrelations, diffusion time as a function of 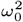 (left) and average number of particles in the focal volume as a function of the focal volume. The fitted corresponding diffusion coefficient is (53 ± 2)*μm^2^/ s* and the particle concentration is (1720 ± 4) /*μm^3^* (averages and standard deviations over 3 measurements of 130 s each). All start-stop (i.e., TCSPC) histograms have 48 ps resolution (bin-with).

Similarly to imaging, we took advantage of the BrightEyes-TTM output signal, which duplicates the central element photon channel, and we fed this signal into the control system of the microscope for a real-time display of the intensity trace and the corresponding autocorrelation function. The autocorrelation curve calculated with the data collected by our BrightEyes-TTM is identical to the curve calculated with the data registered by our National Instruments DAQ reference system (Suppl. Fig. S5): this comparison further validates the proposed TTM.

## Discussion

In this work, we presented an open-source multi-channel time-tagging module specifically designed for fluorescence laser scanning microscopy. We validated and demonstrated the versatility and flexibility of this device by implementing FLISM and FLCS but the module is compatible with any SP-LSM technique: a family of methods which leverage the possibility of new array detectors to analyse the specimen’s fluorescence photon-by-photon.

To the best of our knowledge, this work also represents the first validation of SPAD array detector for combining CCA with fluorescence lifetime analysis. This combination can open to new perspectives for single-molecule fluorescence resonance energy transfer (smFRET) in live-cell (19).

We implemented the TTM architecture on a FPGA, mounted on a readily available development kit, thus providing design flexibility and upgradability, and also allowing fast prototyping and testing of new potential BrightEyes-TTM release versions. For example, the current frame-like data-structure adopted to transfer the data to the PC can be improved in terms of throughput. Indeed, the current data-transfer protocol uses a data-structure which allocates space for all photon channels independently on the effective collection of photons. However, for most of the typical fluorescence applications, it is unlikely of having multiple photon events on the same data-structure. Furthermore, the size of the data that is transferred from the TTM to the PC could be significantly reduced by performing part of the data analysis and the reconstruction directly on the TTM, by leveraging a novel class of DAQ cards equipped with ARM-based processors.

The principal aim of this work is to give to any microscopy laboratory the possibility to implement and further develop single-photon microscopy. The second aim is to trigger the interest of the microscopy community, and establish the BrigthEyes-TTM as a new standard for single-photon LSM experiments. We are fully aware that these aims can be achieved only if the microscopy makers community (20) has full access to this device, thus we releases the results of this work as open-source.

By implementing different low precision reference signals, the TTM can collect photons in synchronisation with many different opto-mechanical devices that could potentially encode in the photon other information, directly on indirectly. For example, polarisation modulators and/or analysers can help tagging photons with an excitation and emission polarisation signature. Acoustic tunable filters and spectrometers can provide an excitation and emission wavelength signature. We envisage a future for which each single fluorescent photon will be tagged with a series of stamps describing not only its spatial and temporal proprieties, as shown in this work, but also its polarisation and wavelength characterisations. A series of new algorithms – based on conventional statistical analysis or machine learning techniques, and data-formats will be developed to analyse and store such new multi-parameter single-photon data-sets. Despite, the Life science community will receive the major benefits from the SP-LSM technology, allowing them to obtain novel insights about the many bio-molecular processes governing Life, we are convinced that SP-LSM will find many applications also in material sciences, in particular in the field of single-dot spectroscopy (21).

## Methods

### Time-Tagging Module Architecture

In the context of SP-LSM, the goal of the TTM is to tag each photon that reaches the SP detector (typically with a maximum count rate of tens of MHz) with a spatial signature and a series of temporal signatures (Suppl. Fig. S6). The spatial signature represents the element of the detector, and thus the spatial position, at which the photon arrives. The temporal signatures are the delays of the photon with respect to specific events: (i) the fluorophore excitation events induced by a pulsed laser; or (ii) the changes of some experimental condition, e.g., the starting of the experiment (absolute time) or the change in position of the probing region. Measuring the photon-arrival time with respect to the excitation event (the so-called start-stop time) typically requires high temporal precision (higher than the SPAD array detectors photon-timing jitters, < 100 ps) and a temporal range larger than the pulse-to-pulse interval of the laser, i.e., 10-1000 ns (i.e., 1-100 MHz pulse frequency). In all other cases, a lower temporal precision (e.g., nanosecond) is needed but the requested temporal range increases up to seconds. To meet these conditions, we implemented different time-to-digital converters (TDCs) for the different types of input signals, i.e., a photon signal, a laser SYNC signal, or a reference (REF) signal. Indeed, the purpose of a TDC is to measure the time interval between two events. In particular, we implemented a fine (picosecond precision) and a coarse (nanosecond precision) TDC

A fine TDC can be implemented either in an application-specific integrated circuit (ASIC) or on an FPGA. Time-to-digital converters implemented by ASIC can provide a temporal precision much higher than FPGA-based TDCs (subpicosecond instead of tens of picoseconds) but the latter offer a shorter development cycle and a lower development cost, and they are easily adaptable to new applications. Because (i) our aim was to develop an upgradable TTM, (ii) most fluorescence applications do not require a sub-picosecond temporal precision, and (iii) SPAD array detectors offer a precision of tens of picoseconds, an FPGA-based TDC was the natural choice.

Within the FPGA-based TDC, the literature reports several different architectures (22). Each architecture offers a compromise between different specifications (e.g., temporal precision, temporal range, temporal resolution, dead-time, linearity, and FPGA resources). By using an FPGA, the delay between two events can be easily measured with a counter, which counts the number of clocks between the two events. The counter approach yields a precision no better than the clock period, which is a few nanoseconds for low-cost devices (as in our case) and – in principle – infinite temporal range. While this precision is suitable for measuring the photon-arrival time with respect to second class events, i.e., a change in the experimental conditions (coarse TDC), they do not satisfy the request for the start-stop time assessment. For the delay with respect to the excitation event, a higher precision (10-100 picosecond, depending by the implementation (22)), is achieved by running the photon signal (i.e., the START signal) through a fast tapped delay line (TDL), and then measuring the position that is reached when the SYNC signal from the laser (i.e., the STOP signal) is received (fine TDC). The downside of the tapped delay line approach is that the maximum delay measurable (i.e., the temporal range) depends on the length of the tapped delay line, and thus on the available FPGA-resources.

To keep low on FPGA resources, and implementing on the same architecture a few coarse TDCs for measuring the second class events and one fine TDC for each SPAD array element, we implemented a sliding-scale interpolating TDC architecture (23), also known as the Nutt method (24) (Suppl. Fig. S7). This method uses a pair of tapped delay lines to measure with tens of picoseconds precision the start-stop time, combined with a free-running coarse counter to extend the temporal range. At the same time, the architecture uses the free-running coarse counter to measure the second class events with a few nanoseconds precision. In particular, the architecture combines *N* +1 (*N* = 21 photon channels in this implementation) tapped delay lines and a coarse counter at 240 MHz to obtain *N* fine TDCs with tens of picoseconds precision (for the start-stop time of each photon channel), and *M* coarse TDCs with a nanosecond precision (*M* = 3 reference channels in this implementation). For both TDCs the temporal range is – in principle – limitless. Notably, each fine TDC uses a dedicated START tapped delay line, but shares with the other fine TDCs the STOP delay line: thus, the start-stop time resulting from each fine TDC is measured with respect to the same SYNC signal, which is what is typically needed in all SP-LSM applications. Furthermore, the high frequency clock (240 MHz in our case) allows keeping the delay-lines short (slightly longer than the clock period, i.e., ~ 4.2 ns), thus reducing both the FPGA-resources needed and the dead-time of the fine TDC. Notably, the dead-time of our TDC (~ 4.2 ns) is shorter than (i) typical SPAD detector hold-off times (> 10 ns) and (ii) typical pulsed laser excitation periods for fluorescence applications (12.5–50 ns).

By using two different tapped delay lines for the START and STOP signals, the architecture ensures that the TDCs are asynchronous with respect to the clock counter. The asynchronous design reduces the non-linearity problem of the FPGA-based TDC and auto-calibrates the tapped delay-lines (Supplementary Note 1, Suppl. Fig S8). Here, we describe a single fine TDC, the parallelization to *N* TDCs is straightforward, and the implementation of the coarse TDC comes naturally with the use of the course counter. Each fine TDC is composed by two flash TDCs, each one containing a tapped delay line and a thermometer to binary converter (Supplementary Note 3, Suppl. Fig. S9). The START delay line measures the difference Δ*t*_START_ between the START signal (photon) and the next active edge of the counter clock. Similarly, the STOP delay line measures the difference Δ*t*_STOP_ between the STOP signal (SYNC) and the next active clock rising-edge. Thanks to the free-running coarse counter the architecture is able also to measure *n*_photon_ and *n*_SYNC_, respectively the number of clock cycles between the start of the experiment and the photon (START) signal or the SYNC (STOP) signal. Given these values, (i) the start-stop time Δ*t* equals to 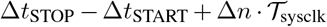, where Δ*n* = *n*_SYNC_ – *n*_photon_, and 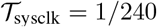 MHz is the clock period; (ii) the *coarse* absolute time 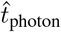, i.e., the nanoseconds precision delay of the photon with respect to the beginning of the experiment, equals to *n*_photon_ · *t*_sysclk_. Similarly, the nanosecond delay from the REF signal 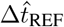 equals to 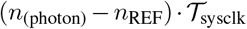, where *n*_REF_ is the number of clock cycle between the beginning of the experiment and the REF signal. The TTM architecture can provide only integer values. In particular, the course counter provides the values *n*_photon_ and *n*_SYNC_, and the time-to-bin converters of the flash TDCs provide the values Δ*T*_photon_ and Δ*T*_SYNC_. After the raw data is transferred to the PC, the calibration is used to calculate the time values Δ*t*_photon_ and Δ*t*_SYNC_ (Supplementary Note 1), and the application-dependent analysis software allows calculating the time interval between specific events (e.g., Δ*t*, 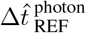). This approach does not preclude applications in which some other notion of an absolute time is required, such as the absolute photon time (e.g., coarse 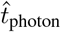, or fine *t*_photon_), rather than the time interval between two events.

The TTM architecture also contains hit-filters (Suppl. Fig. S10), an event-filter (Suppl. Fig. S11), and a data-module in charge of preparing and transferring the data to the PC. The hit-filter is a circuit used to shape and stabilise the input signal. When a hit (photon or SYNC event) occurs, this circuit keeps the signal high until it gets sampled by the internal clock. The hit-filter guarantees the stability of the TDL input signal until the next rising edge event of the internal FPGA clock. The event-filter is a data-input/output filter that reduces the data throughput rate by avoiding to transmit information when no photons are detected. The data-module are the data following a well-defined structure, and buffers it into a FIFO, before being transferred to the PC. In its current implementation, data-transfer is done with a USB 3.0 protocol, but it is compatible with other data communication protocols, such as a PCIe.

We implemented the BrigthEyes-TTM architecture on a commercially available Kintex-7 FPGA evaluation board (KC705 Evaluation Board, Xilinx), featuring a state-of-the-art FPGA (Kintex-7 XC7K325T-2FFG900C), and – very important for the versatility of the architecture, a series of hardware components (e.g., serial connectors, expansion connectors, and memories). In particular, the different serial connectors potentially allow for different data-transfer rates, and the expansion connectors allow compatibility with other detectors and microscope components. Notably, the full TTM system currently implemented uses 75% of the FPGA resources (Suppl. Fig. S12), thus allowing for the introduction of new photon or reference channels or other functions.

The TTM design here presented uses, to transmit the data to the PC via USB 3.0, a FX3-based board (Cypress Super-Speed Explorer kit board, CYUSB3KIT) connected through an adapter card (CYUSB3ACC) to LPC-FMC connector of the Kintex-7 evaluation board. In order to use the FX3, a dedicated module in the FPGA was developed. It has a simple interface (essentially FIFO with a data-valid flag) for the data transmission and it manages the FX3 control signals and the data-bus. The module was designed to work with the FX3 programmed with the SF_streamIN firmware part of the AN65974 example provided by Cypress.

### Test-Bench Architecture and Analysis

We performed the code-density test (Supplementary Note 2) and the single-photon precision measurement by connecting the BrightEyes-TTM to a dedicated signal generator, named SYLAP (Suppl. Fig. S13). We developed SYLAP for this specific project, however, it can help any application which requires synchronised signals with tunable delay. We implemented SYLAP on an FPGA-evaluation board identical to the one used for the BrigthEyes-TTM (i.e., KC705 Evaluation Board, Xilinx). The SYLAP architecture generates a fixed frequency clock, which we used to simulated the laser SYNC signal, and a synchronised pulse train, which we used to simulate the photon signal. Key features of the SYLAP are (i) the possibility to adjust the delay from the clock and the pulse with a granularity of 39.0625 ps; (ii) the possibility to set the clock period and the pulse duration with a granularity of 2.5 ns. The native USB 2.0 serial port of the evaluation board allows configuring the SYLAP. The timing jitter between the clock and the pulse is 13 ps, measured with Keysight DSO9404A 4 GHz bandwidth oscilloscope.

Since both the BrightEyes-TTM and the SYLAP signal generator use the same type of board, we also implemented both systems on the very same FPGA (thus same board). Importantly, in this internal implementation of SYLAP we separated the clock-domains and the clock sources of the two projects. We performed a single-photon precision experiment to compare the stand-alone SYLAP configuration (i.e, two different boards connected with coaxial cables), with the internal configuration, and we did not observe substantial differences. The SYLAP internal implementation demonstrates the flexibility of our TTM architecture in integrating new features.

To perform the code-density test (Supplementary Note 2) we used the clock signal from SYLAP (as SYNC signal) and as random photon signals (i.e., temporally correlated), we used the TTL signal from an avalanche photo diode (APD, SPCM-AQRH-13-FC, Perkin Elmer). We illuminated the APD with natural light, maintaining a photon-flux well below the saturation value of the detector.

To perform the single-photon precision experiment for all photon channels of the TTM, we took again advantage of the flexibility of our TTM architecture, and in particular the possibility to reprogram the FPGA according to different requests. Indeed, we implemented physical switches which allow simultaneously connecting a single input channel of the board to all 21 photon channels. This feature gives the possibility to measure the same event with all photon channels.

### Single-Photon Microscope

#### Optical Architecture

To test the BrigthEyes-TTM we used a custom-built SP laser-scanning microscope (Suppl. Fig. S14) previously implemented to demonstrate confocal-based FFS with SPAD array detectors (7). In short, the microscope uses a 485 nm pulsed laser (LDH-D-C-485 PicoQuant, driven by a PicoQuant PDL 828 Sepia II Multichannel Picosecond Diode Laser driver) for excitation with a tunable repetition rate *f* (in this work we used 5, 10, 20, 40, 80 MHz). After passing through a 488/10 nm clean-up filter, the laser light was reflected by a dichroic beam splitter (ZT405/488/561/640 rpc, Chroma Technology Corporation) towards the galvanometric scanning mirrors. The scanning system was coupled to a 50 mm Leica scan lens and a 200 mm Leica tube lens. We performed all measurements with a 100×/1.4 Leica objective lens. Focusing, axial scan, and scouting of the sample region-of-interest, were performed with a 3-axis stage (Nano-LP Series, Mad City Labs). The fluorescence signal was collected in de-scanned mode, in which it passed through the dichroic beam splitter, a 488 nm long pass filter and a fluorescence emission filter (ET550/25, Chroma Technology Corporation). A 250 mm lens conjugated with the scan lens was installed to obtain a 1.5 Airy unit field of view on the SPAD array detector, which meant that the detector also acted as a pinhole to remove the out-of-focus fluorescence background. The measurements were performed with a 5× 5 SPAD array detector fabricated using BCD technology (16). The hold-off time was set to 100 ns.

#### Control System

To control the SP laser-scanning microscope, we built a LabVIEW system inspired by the Carma control system (15, 25). The LabVIEW software uses an FPGA-based National Instrument (NI) data-acquisition-card (NI USB-7856R; National Instruments) to control all microscope instruments, such as the galvano-metric mirrors, the 3-axis piezo stage, and the SPAD array detector (through an initialisation protocol). The pixel, line, and frame clocks were delivered as digital outputs. Finally, the control system was able to record the photon signal from the SPAD array detector and transfer the data to the PC (via USB 2.0), where the LabVIEW software allowed real-time visualisation of the images – in the case of imaging – or the intensity time-traces – in the case of FFS. Notably, when the SP laserscanning microscope was used in time-tagging modality, the photons-signals were delivered to the BrightEyes-TTM. In this case, the BrightEyes-TTM read the photon-signal thanks to a custom I/0s daughter card connected via the FPGA mezzanine connector (FMC). The same card was used to deliver the duplicate signal of the central element of the SPAD array to the LabVIEW control system. The pixel, line, and frame signal passed through a custom-made buffer to match the impedance between the NI and the Xilinx FPGA card. The SYNC signal provided by the laser driver was converted from NIM to TTL (NIM2TTL Converter, MPD) and read by the Xilinx DAQ cards via a SMA digital input. The data recorded by the BrightEyes-TTM were transferred to a PC via USB 3.0. To compare the TTM with a commercial reference system, the SYNC (NIM) signal and the signal from the central element of the SPAD array could also be sent to a commercial TTM card (DPC-230, Becker & Hickl GmbH), running on a dedicated PC.

### Data Transfer

#### The data structure

To transfer the data from the TTM to the PC, we designed a simple data protocol, whose major advantage is the scalability to add photon channels, as well as its flexibility. In a nutshell, the protocol perceives the SP detector array as a fast camera with a maximum frame-rate of 240 MHz, i.e., the frequency at which the whole TTM architecture works. Under this scenario, the data protocol foresees a unique frame-like data structure streamed to the communication port (in this implementation the USB 3.0) in 32-bits long words (Suppl. Fig. S15). Since the current Bright-Eyes TTM version can read 21 photon channels, it needs to transmit one header word and seven payload words (3 photon channels *per* payload word), thus 256 bits in total. To check the data receiving order and to avoid misinterpretation, each word includes 4 bits as identifier (ID). The first word (header, ID = 0) includes the following information: (i) three bits used as boolean-flag for the reference (REF) events (in our applications the pixel, line, and frame clocks); (ii) the 8 bits representing the value measured by the tap-delay line of the SYNC channel Δ*T*_STOP_ (in our application the synchronisation signal from the pulsed laser), and (iii) the SYNC data valid boolean-flag which confirms that a SYNC event has occurred; (iv) the 16 bits representing the number of clock cycles of the free-running 240 MHz counter. The successive payload words (ID > 0) contain the information of three photon channels. For each photon channel the word contains: (i) the 8 bits representing the value measured by the tap-delay line of the respective photon channel Δ*T*_START_(*ch*); (ii) the channel data-valid boolean-flag which confirms that an event in that channel has occurred. Each payload word contains also a general purpose single bit, which can be used to implement another REF_N_ signal.

Since the data structure is 256 bit long, and, in principle, we have to transmit the structure with a rate of 240 MHz, the data throughput would be 7.68 Gbps. However, the USB 3.0 has an effective bandwidth of 3 Gbps. For this reason, the data structure is transmitted only if one of the following conditions occurs: (i) a START event in one of the photon channels, e.g., a photon event; (ii) a STOP event following a START event, e.g., a laser SYNC event occurs after a photon event (event filter); (iii) a REF event, e.g., a pixel/line/frame event; (iv) a force-write event. The force-write event is fundamental for reconstructing any time measurement (relative or absolute) that requires the coarse counter values. Indeed, since the data structure uses only 16 bit for storing the 240 MHz course counter value, the counter needs to reset every 273 *μs*. To guarantee the possibility to always reconstruct the relative or absolute value for each event, we implemented an internal trigger which forces to transmit the data structure at least once every 17 *μs*, value that corresponds to one sixteenth of the course counter reset period.

To improve the robustness of the data transfer process, we mitigated the data throughput peaks by buffering the data in a large (512 kB) BRAM-FIFO on the same FPGA. The TTM code contains a mechanism that guarantees that in case the FIFO gets filled over a certain threshold, i.e., the average data throughput exceeds the USB 3.0 data bandwidth (because of PC latency, a high rate of events, or other reasons), the TTM enters temporarily in a fail-safe mode, giving priority to some type of events. For example, in the case of imaging, the TTM gives priority to the pixel/line/frame flags and to the force-write events. This strategy guarantees the reconstruction of the absolute time of each event (photon or REF) and the image.

In current TTM-implementation, aUSB 3.0 effectively transfer the data to the PC. Here, a data receive software checks the data integrity of the data structures received (i.e. the IDs must be in correct order), and, if the data structure is properly received, it sequentially write the data to the raw file without any process (raw data). The data receiver uses the *libusb-1.0*, and developed in C programming language.

#### The pre-processing

To create a user-friendly dataset (Suppl. Fig. S16), we pre-processed the raw dataset. Since the free-running counter has 16 bits, it resets every 273 us; therefore, to obtain a consistent monotonic counter *n*, we update the 16 bits long free-counter provided by the data structure: when the free-counter value is lower than the one in the previous data structure, the value 2^16^ is added to it and added to the following counter values.

With the monotonic counter *n*, it is possible to calculate the arrival time of each photon event with respect to a REF event or with respect to the SYNC event (start-stop time). While the former calculation is trivial, the latter is more complex. Since the TTM architecture uses a start-stop reverse strategy to reconstruct the photon start-stop time – for a given channel *ch*, it is necessary to recover the information about successive SYNC laser events. Notably, this information can be contained in the same data structure of the photon, or in a successive data-structure. The pre-processing step identifies for each STOP (SYNC) event the corresponding START (photon) event and creates a table. Each table row contains a STOP (SYNC) event and includes: (i) the relative monotonic counter *n*_SYNC_; (ii) the relative TDL value Δ*T*_STOP_; an entry for each photon event linked to this specific STOP (SYNC) event. In particular, each entry contains: (i) the number of elapsed clock cycles Δ*n*(*ch*) = *n*_SYNC_ – *n*_photon_(*ch*); (ii) the TDL value Δ*T*_START_(*ch*).

To simplify the reconstruction of the image, in this phase we also pre-processed the REF information for the pixel/line/frame. We used the pixel/line/frame events to include in the table the columns *x*, *y* and *fr.* Namely, the pixel event increases the *x* counter; in case of the line event, the *x* counter resets and the line counter *y* increases; in case of the frame event, both the *x*, *y* counters reset, and the *fr* counter increases.

We saved the pre-processed dataset in a single compressed HDF5 file composed by different tables: the main table and the 21 tables referred to as the photon channels. All tables use a unique column identifier (*idx*), which allows the applications software (e.g., FLISM software, FLCS software) to easily merge the information. The main table has a row for each SYNC channel event. Each row contains the corrected monotonic counter *n*_SYNC_, the coordinates *x*, *y*, *fr*, the TDL value Δ*T*_STOP_ value, and the unique row index *idx*. The photon channel tables have a row for each photon. Each row contains the TDL value Δ*T*_START_(*ch*), the elapsed clock cycle Δ*n*(*ch*), and the index *idx* of the row of the corresponding sync event.

#### Calibration

Notably, the HDF5 file contains all the information as received by the TTM, but structured in a way that is much easier to access. However, this information still contains numbers of cycles or tapped-delays. A calibration offline phase allows transforming this information in temporal information. In particular, the calibration (Supplementary Note 1, Suppl. Fig S17) transforms each TDL value Δ*T*_START_(*ch*) or Δ*T*_SYNC_ in a temporal value Δ*T*_START_(*ch*) or Δ*t*_SYNC_, which we can use to calculate all (both relative and absolute) temporal signatures of each event. The output of the calibration is again an HDF5 file, with a structure similar to the uncalibrated file. The main table has a row for each SYNC channel event. Each row contains the absolute SYNC time 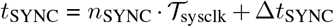, the coordinates *x*, *y*, *fr* and a unique row index *idx*. The photon channel tables have for each row the start-stop time Δ*t*(*ch*), and the index *idx* of the row of the corresponding SYNC event.

#### Applications

Depending on the application, we further processed the calibrated HDF5. In general, any application requires the generation of the start-stop time (or TCSPC) histogram, which is simple the histogram of a series of Δ*t* values. Because, the auto-calibration steps (Supplementary Note 1), the Δ*t* values are float values, thus the bin-with of the histogram (i.e., the temporal resolution) can be choose almost arbitrary by the user. In all this work we used 48 ps, which is a value well below the system IRF (200 ps, Fig. 1) and in the range of the average delay value of the elements (i.e., CARRY4) used to build the tapped delay lines (Supplementary Note 1. In the case of FLISM analysis, the calibrated data are binned into a multidimensional intensity (photon-counting) array (*ch,x,y, fr,τ*). The *τ* dimension is the start-stop histogram for the given other coordinates. This multidimensional array can be saved in HDF5 and processed with *ad hoc* scripts or software like ImageJ. The other dimensions *ch*, *x*, *y*, *fr* are self-explainable. In the case of FFLS analysis, we ignored the *x*, *y* and *fr* information of the calibrated HDF5 and we did not apply any further processing. Indeed, the FFLS needs a list of photon events for each channel, in which each photon has an absolute time – with respect to the beginning of the experiment, and the start-stop time (Δ*t*). By using *t*_SYNC_ as absolute time (instead of *t*_photon_ the calibrated HDF5 file contains already all information structured in the best way.

### FLISM and FLFS Analysis

#### FLISM Reconstruction and Analysis

We reconstructed the ISM images by using the adaptive pixel-reassignment method (15). In short, we integrated the 4D dataset (*ch,x,y,τ*) along the *τ* dimension, we applied a phase-correlation registration to register all the *ch* images with respect to central image, i.e., *ch* = 10. The registration generated the so called shift-vector fingerprint (*s_x_*(*ch*), *s_y_* (*ch*)). To obtain the ISM intensity-based images, we integrated along the *ch* dimension the shifted dataset. To obtain the lifetime-based ISM image, we start from the 4D dataset (*ch,x,y,τ*), for each *τ* value we used the same shift-vector fingerprint to shift the relative 2D image, and we integrated the result along the *ch* dimension (Supl. Fig. S18). Finally, we used the resulting 3D dataset (*x,y,τ*) and the FLIMJ software (26) to obtain the *τ_fl_* map (singleexponential decay model) and the FLISM image. Alternatively, we applied the phasor analysis on the same 3D dataset. We calculated the phasor-plots pixels coordinates (*g, s*) using cosine and sine summations (27) (28). To avoid artefacts in the phasor plot calculation we did the MOD mathematical operation of the TCSPC histograms with the laser repetition period value (28). Phasor-plot analysis measurements were referenced to a fluorescein solution which was used to phasor-calibrate the entire acquisition system and thus accounting for the instrument response function of the complete setup (microscope, detector and TTM).

To demonstrate the resolution enhancement achieved by ISM, we performed a Fourier ring correlation (FRC) analysis (29, 30). The FRC analysis require two “identical” images, but obtained from two different measurements in order to contain a different noise realisations. The two images are correlated to obtain the effective cut-of-frequency (i.e., the frequency of the specimen above the noise level) of the images, and thus the effective resolution. Since, in this work build-up the images photon-by-photon, we can used the temporal tags of the photons, to generate simultaneously two 4D data-sets (*ch,x,y,fr,τ*) required by the ISM analysis. In particular, we odd-even sort each photon in the two images by using the Δ*T*_START_ and Δ*T*_START_ integer values. As explained for the sliding-scale approach, the photons distribute uniformly across the START and STOP tapped-delay lines, thus the method generates two statistical independent dataset with similar photons counts.

#### Correlation Calculation and Analysis

We calculated the correlations directly on the lists of the photon absolute time (31). For the sum 3×3 and sum 5 × 5 analysis, the lists of all the relevant SPAD channels were merged and sorted. Then, the data was split into chunks of 10 s and for each chunk the correlation was calculated. The individual correlation curves were visually inspected and all curves without artefacts were averaged. To obtain the filtered correlation curves, the same procedure was followed, except that a weight was assigned to each photon based on its start-stop time (32, 33). The weights were obtained from the start-stop time histograms of each channel. Only photons in time bins between the peak of the histogram and about 10 ns later were included. In this way, photons that are likely to not originate from the sample were removed. First, the TCSPC histogram of each channel was fitted with a single exponential decay function *H*(*t*) = *A* exp(-*t/τ_fl_*) + *B*, with the amplitude *A*, lifetime *τ_fl_*, and offset *B* as free fit parameters. The filters were calculated assuming a single exponential decay with amplitude 1 and lifetime τ*_fl_* for the fluorescence histogram and a uniform distribution with value *B/A* for the afterpulse component. Then, for each photon, a weight was assigned equal to the value of the fluorescence filter function at the corresponding photon start-stop time. E.g., photons that were detected directly after the laser pulse were assigned a higher weight than photons detected some time later, since the probability of a photon being a fluorescence photon decreases with increasing start-stop time. Correlations were calculated taking into account these photon weights. The second filter function was not used for further analysis, since it would amplify the afterpulse component and attenuate the fluorescence contribution. All correlations were fitted with a 1-component model assuming a Gaussian detection volume (i.e., probing region). For the circular FCS measurements (34), the periodicity and radius of the scan movement were kept fixed while the amplitude, diffusion time, and PSF size were fitted. For the conventional FCS measurements, the PSF size was kept fixed at the values found with the circular scanning FCS, and the amplitude and diffusion time were fitted.

### Samples Preparations

For TTM characterisation and validation, we used Fluorescein (46955, free acid, Sigma-Aldrich) and potassium iodide (60399-100G-F, BioUltra, >= 99.5% (AT), Sigma-Aldrich). We dissolved Fluorescein from powder into DMSO and then we further diluted to a 1:1000 v/ concentration by adding ultrapure water. For the Fluorescein quenching experiments we diluted the 1:1000 Fluorescein solution at different ratio with the potassium iodide (KI) quencher (from 1:2 to 1:256). All samples were made at room temperature. A fresh sample solution was prepared for each measurement. For imaging experiments we used: (i) a sample of 100 nm fluorescent beads (yellow-green FluoSpheres Q7 Carboxylate-Modified Microspheres, F8803; Invitrogen). We treated the glass coverslips with poly-L-lysine (P8920; Sigma-Aldrich) for 20 min at room temperature, and the we diluted the beads in Milli-Q water by 1:10,000 v/v. We drop-casted the beads onto the coverslips, and, after 10 min, we washed the coverslips with Milli-Q water, we dried under nitrogen flow, and we mounted overnight with Invitrogen ProLong Diamond Antifade Mounting Medium (P36965); (ii) a prepared microscope slide (IG-4011 Imaging Set, Abberior) containing fixed mammalian cells in which vimentin were stained with the STAR-GREEN dye; (iii) a fresh prepared microscope slide containing fixed Hela cells stained for tubulin visualisation. We cultured Hela cells in Dulbecco’s Modified Eagle Medium (Gibco, ThermoFisher Scientific, Wilmington) supplemented with 10% fetal bovine serum (Sigma-Aldrich) and 1% penicillin/streptomycin (Sigma-Aldrich) at 37 °C in 5% CO2. One day before immunostaining, we seeded Hela cells onto coverslips in a 12-well plate (Corning Inc.). We incubated cells in a solution of 0.3% Triton X-100 (Sigma-Aldrich) and 0.1% glutaraldehyde (Sigma-Aldrich) in the BRB80 buffer (80 mM Pipes, 1 mM EGTA, 4 mM MgCl, pH 6.8, Sigma-Aldrich) for 1 min. We fixed Hela cells with a solution of 4% paraformaldehyde (Sigma-Aldrich) and 4% sucrose (Sigma-Aldrich) in the BRB80 buffer for 10 min and then we washed three times for 15 min in phosphate-buffered saline (PBS, Gibco™, ThermoFisher). Next, we treated cells with a solution of 0.25% Triton-X100 in the blocking buffer for 10 min and we washed three times for 15 min in PBS. After 1 h in blocking buffer, a solution of 3% bovine serum albumin (BSA, Sigma-Aldrich) in BRB80 buffer, we incubated the cells with monoclonal mouse anti-*α*-tubulin antibody (Sigma-Aldrich) diluted in the blocking buffer (1:1000) for 1 h at room temperature. The alpha-tubulin goat antimouse antibody was revealed with Alexa Fluor 488 goat antimouse (Invitrogen, ThermoFisher Scientific). We rinsed Hela cells three times in PBS for 15 min. Finally, we mounted the cover slips onto microscope slides (Avantor, VWR International) with ProLong Diamond Antifade Mountant (Invitrogen, ThermoFisher Scientific). For fluorescence fluctuation spectroscopy experiments we used: (i) YG carboxylate fluo-Spheres (REF F8787, 2% solids, 20 nm diameter, actual size 27 nm, exc./em. 505/515 nm,Invitrogen, ThermoFisher) diluted 5000× in ultrapure water. A droplet was poured onto a cover slip for the FLFS measurements; (ii) The goat anti-Mouse IgG secondary antibody with Alexa Fluor 488 sample (REF A11029, Invitrogen, ThermoFisher) was diluted 100× in PBS to a final concentration of 20 *μ*g/mL. 200 *μL* of the resulting dilution was poured into an 8 well chamber previously treated with a 1% BSA solution to prevent the sample to stick to the glass. All samples were made at room temperature. A fresh sample solution was prepared for each measurement.

## Data availability

As keen proponents of open science and open data, we have made the raw time-tagged data, which supports the findings of this study, publicly available from Zenodo, https://doi.org/10.5281/zenodo.4912656. Full build instructions for the BrightEyes-TTM is available through our GitHub repository https://github.com/VicidominiLab/BrightEyes-TTM.

## Code availability

The firmware and the VHDL/Verilog source code for implementing time-tagging on the FPGA evaluation board, the data-receiver software to install on the personal computer, and the operational software with are accessible through the Vicidomini Lab Github https://github.com/VicidominiLab/BrightEyes-TTM.

## ACKNOWLEDGEMENTS

This research was supported by Fondazione San Paolo, “Observation of biomolecular processes in live-cell with nanocamera”, No. EPFD0098 (E.S. and G.V.) and by the European Research Council, Bright Eyes, No. 818699 (G.T. and G.V.). We thank Prof. Alberto Diaspro and Dr. Paolo Biancchini (Nanoscopy & NIC@IIT, Istituto Italiano di Tecnologia) for usefull discussion; Dr. Michele Oneto (Nikon Imaging Center) for support on the experiments; Alessandro Barcellona for design and implementation of the custom-made buffer (Electronic Design Laboratory, Istituto Italiano di Tecnologia); Sabrina Zappone (Molecular Microscopy and Spectroscopy, Istituto Italiano di Tecnologia) for cell preparation and labelling; Prof. Alberto Tosi, Prof. Federica Villa, Dr. Mauro Buttafava (Politecnico di Milano), Dr. Marco Castello, and Dr. Simonluca Piazza (Istituto Italiano di Tecnologia and Genoa Instruments) for useful initial discussions in the time-to-digital design and for the realisation of the single-photon-avalanche-diode detector array.

## AUTHOR CONTRIBUTIONS

G.V. conceived the idea and designed the research; G.V. and M.C. supervised the research, A.R., M.D., and F.D. wrote the firmware for the time-tagging module and the data-receiver; M.D. with support from A.R. wrote the operational software; A.R., E.S., M.D., S.K., and G.T. wrote the imaging analysis software; E.S., M.D., and L.L. wrote the FFS analysis software; A.R. and M.D., characterised and validate the TTM module with the test-bench architecture; E.S. build the single-photon microscopy; E.S. and E.P. with support from A.R. performed the single-photon microscopy experiments; A.R., E.S., M.D., E.P., S.K., G.T., and G.V., analysed the data. A.R., M.D. and G.V., build the GitHub repository; G.V., with supports from A.R., E.L., and M.D., wrote the manuscript with edits from all authors.

## Supplementary Note 1: Sliding-Scale Technique and Auto-Calibration

The sliding scale approach is the most suitable FPGA-based TDC architecture for the BrightEyes-TTM because it incorporates features that are crucial for single-photon microscopy applications. In this supplementary section, we explain how the sliding-scale design allows for a flexible multi-channel TDC implementation, which reduces the well-known non-linearity problem of FPGA-based TDCs and saves up on FPGA resources, allowing the BrightEyes-TTM to be used with different laser sync rates and allowing auto-calibration of the TTM.

The sliding-scale technique uses multiple flash TDC modules, one for each implemented channel (START channels), and a dedicated one for the STOP channel (i.e. the laser SYNC signal), deployed in the FPGA together with a coarse free-running digital counter (Suppl. Fig. S7). A key feature of this technique is that the flash TDC readouts and the coarse counter are synchronous with the FPGA internal reference clock, which is totally uncorrelated with respect to the STOP signal and therefore also to the photon arrival time. The sliding-scale technique can thus be defined as an asynchronous design (1–3). An advantage of this design is that all the input signals (STARTN and STOPSYNC) produce time-tagging values over the full available dynamic range of a flash TDC. In short, the very same experiment can be used for calibration (e.g., estimate for each delay unit of the flash TDC module the effective duration, Suppl. Note 3) independently of all flash TDCs, and to further compensate for the non-linearity.

### Sliding scale technique

To tackle the linearity issue, the fundamental sliding scale TDC architecture is made up of a flash TDC module to sample START events (i.e. incoming photons) and a second flash TDC module to acquire the STOP signal (i.e. the laser sync signal) (Suppl. Fig S8). As described in literature (4), a conventional FPGA-based flash TDC should not be used because of its lack of linearity across the probed temporal range, caused by intrinsic non-linearities in the FPGA fabric. The non-linear behaviour of flash TDC modules is shown in Figure S8 (a). In the sliding scale technique, instead of using a single flash TDC in which the delays between the STARTs and STOP signals are directly measured, two independent flash TDC modules are used in combination with a coarse counter. One flash TDC measures Δ*t*_photon_, i.e., the arrival time of the photon (START) with respect to an internal FPGA reference clock. The second TDC measures Δ*t*_SYNC_, i.e., the delay of the laser sync (STOP) with respect to the same FPGA reference clock, Fig S8 (a). The coarse counter is needed to calculate the time difference (Δ*t*) between the photon and the laser pulse, Eq. S1 (see also Fig S8 a):

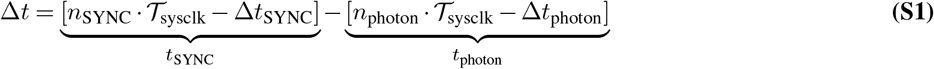

Since the photon and laser sync signal are asynchronous with respect to the FPGA clock, the same time interval Δ*t* is measured in different regions of the flash TDC module for different photons. Using the sliding scale approach, identical time intervals will therefore yield slightly different flash TDC responses. However, these non-linearities are averaged out over multiple observations (Suppl. Fig. S8 b). In order to save up on FPGA resources (to allow to further scale up the current design) and to take advantage of the TTM auto-calibration system described below, the calculations and time conversions for computing Δ*t*, Δ*t*_photon_ and Δ*t*_SYNC_ take place on a host computing unit in a post-processing phase. Here, the flash-TDC readings and the registered coarse counter values are merged for each channel to reconstruct the history of each photon (Suppl. Fig. S15). In the proposed TTM architecture, the conjunction of separate flash TDC modules and a common 16 bit-wide coarse free-running counter for all the implemented channels not only levels out the non-linearity but also offers the possibility to (i) keep track of the experiment time, allowing the TTM to have a virtually infinite temporal range, and (ii) to deal with different laser sync rates. Together with the photons and laser sync arrival times, the wrapping events of the coarse counter are also registered and used to calculate the experiment time by simply combining the number of wraps with the period of the FPGA reference clock (240 MHz i.e. ~4.2 ns). The ability of the TTM to reconstruct the time axis based on the internal FPGA clock also allows the TTM to be used with different laser sync frequencies without needing to change the FPGA firmware. As a result, the TTM can be used to work at different temporal ranges, having always the same linear response.

### Auto-calibration

Thanks to the sliding-scale approach, the TTM calibration procedure can be carried out using the same experiment data. In order to calibrate a flash TDC module (i.e. assessing the time width of all the small delay elements that constitute a flash TDC, Suppl. Note 3) a statistical code density test needs to be performed (Suppl. Note 2). By sampling and accumulating random photon events, one can build up a look-up-table (LUT) that contains the time-conversion factors of all the possible TDC readouts to time-tag the incoming photons. In the sliding-scale approach, all the input signals (photons and laser sync events) are, by default, unsynchronized with the FPGA clock. The statistical code density test is thus built-in into the tagging architecture mechanism design: the TTM can be calibrated with experiment data, hence the name auto-calibration. All the deployed flash TDC modules are calibrated bin-by-bin in a post-processing phase using the reconstructed experiment data. In this way, voltage and temperature changes will not affect the data and timing reconstruction. In conclusion, although the auto-calibration is conducted off-line (i.e., not in real-time), it has the advantage that no ad-hoc calibration acquisitions are needed and, since it can be implemented on the computer, it does not need extra resources on the FPGA chip.

### Bin-by-bin calibration

In the sliding scale approach, the flash TDC clock (FPGA clock) is not correlated with the START_n_ or STOP inputs, i.e., all measured inputs have the same probability of falling into any bin of the tapped delay line of the flash TDC module (Suppl. Fig. S9 a). Since the bin widths (i.e. the delay element values of the tapped delay line) are not all equal due to intrinsic FPGA fabric irregularities, a photon or sync hit is more likely to fall into a wider bin than into a narrower bin (4). In order to calibrate the time response of the TTM, the histograms of the hit counts as a function of the arrival bin of each deployed flash TDC module are used. After having collected a large number of START_n_ or STOP events from a measurement, the cumulative event count in each bin is proportional to its width. For example, if a total of N hits are accumulated into the histogram (Suppl. Fig. S17 a), assuming these hits are evenly spread over ~4.2 ns, which is the period of 240 MHz FPGA clock driving the flash TDC, then the width of an N_i_-count bin is w_i_ = N_i_*(4200 ps)/(N) (Suppl. Fig. S17 b). In the bin-by-bin calibration procedure, the widths wi of all tapped delay line CARRY elements (Suppl. Note 3) are measured and stored in an array w_k_, then the calibrated time responses Δt_i_, corresponding to the center of i-th bin, can be calculated according to Eq. S2 below (5, 6):

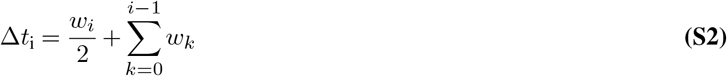

In this way, all the different Δti time contributions of all the flash TDC module delay elements, can be assessed and used to correctly time-tag both the photons (START_n_) and the laser sync line (STOP) (Suppl. Fig. S17 c).

### TCSPC histogram binning

Since we use the sliding-scale (or Nut) method to implement the fine TDC, and an off-line (postacquisition) bin-by-bin calibration, the user can arbitrary choose the bin width of the start-stop (TCSPC) histogram. In this work we always used the same bin width value, i.e., 48 ps. We used this value for the code density test (i.e., least-significant-bit (LSB) is equal to 48 ps), for the single-shot precision measurements, and for all the experimental measurements. This value is much lower than the system IRF full-width-at-half maximum (200 ps), and is in the same range of the CARRY’s average temporal length, i.e., the length of the tapped delay elements used to implement the TDC. Supplementary Figure S19 show the calculated (bin-by-bin calibrated) wk bin widths for all the N = 21 deployed flash-TDC modules in a typical code-density experiments. The average values for the bin widths is 42.98 ± 25.27 ps. Notably, this information is not used in any TCSPC histogram reconstruction, since the bin width is arbitrary choose by the user. Supplementary Figure S20 show the very same start-stop histogram (i.e., the same experiment) for different bin-width values. The histogram change substantially shape only for bin-width much higher than the with of the system IRF. This value is much lower than the system IRF full-width-at-half maximum (200 ps) and in the same range of the CARRY’s temporal length, i.e., the length of the tapped delay element.

The average of the calculated (bin-by-bin calibrated) w_k_ bin widths for all the N = 21 deployed flash-TDC modules was 48ps. For this reason, 48 ps was chosen as the least-significant-bit (LSB) value and used as the standard bin width to make histograms of the photon counts. Since we use the Nutt-sliding scale module and off-line post-acquisition calibration, the bin width can be virtually chosen to be any value greater than 48 ps (Suppl. Fig. S20).

## Supplementary Note 2: Statistical Code Density Test

In this supplementary section, we explain the statistical code density test used to characterize the linearity of the BrightEyes-TTM. The code density test allows measuring (i) the time response of each time-bin of the TDC and (ii) the deviation of TTM measurement readouts from the actual time of arrival of a photon. One of the most important parameters to analyse in order to asses the system response of a time-tagging device is the system linearity. A statistical code density test consist of feeding the TTM random (uncorrelated) photons with respect to the sync reference signal. Here, we generated random photons by connecting an avalanche photodiode (APD) set up in a dark room to the TTM (Suppl. Fig S13). After having reconstructed the histogram *H*_(*i*)_ of the collected random events (using bin-by-bin auto-calibrated data), *H*_(*i*)_ histogram data was used to compute two benchmark indicators of a system’s linearity: the differential non-linearity (DNL) and the integral non-linearity (INL) (Suppl. Fig 1 a). The DNL and INL may seem counter-intuitive indices as the DNL describes the non-linearity amongst the different time-bins of *H*_(*i*)_ (i.e., how much all the bin widths in the temporal range differ from each other) and the INL to what extent the system is non-linear (i.e., to what extent the system is capable of precisely measuring time with respect to an ideal time-tagging device). The relevant values are the standard deviations of the DNL and INL, which are expressed in LSB. In agreement with the Xilinx Kintex-7 XC7K325T-2FFG900C datasheet, an LSB of 48 ps, corresponding to the time delay associated with the coarseness of the CARRY4 element (employed as the fundamental unit in the flash TDC module) was used to reconstruct *H*_(*i*)_ and compute *σ*_DNL_ and *σ*_INL_ (7).

### DNL - Differential non-linearity

When performing a statistical code density test, the reconstructed time histogram *H*_(*i*)_ should ideally be a constant flat line, indicating that every time-bin has an equal time-width within the measured temporal range. In reality, due to intrinsic FPGA fabric inconsistencies, *H*_(*i*)_ shows a ripple that needs to be characterised in order to maximise the timing precision and accuracy. The aim of computing the DNL is to understand to which degree the difference in time widths of all the possible time bins (which is the cause of the ripple) deviates from a common average value. In other words, the DNL is used to understand the relative contribution of all the time bins when reconstructing times of arrival of photons in the TDC temporal range. The DNL is calculated according to Eq. S3:

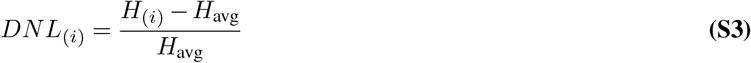

Here, *H*_(*i*)_ is the reconstructed histogram of the dark counts and *H*_avg_ its average value. *DNL*_(*i*)_ represents the deviation of the i-th time bin from the *H*_avg_ value: the lower this deviation, the flatter the *DNL*_(*i*)_ plot will be. If the *DNL* is constant, all time-bins have the same width and the system is perfectly linear. Having set the LSB to 48 ps, *σ*_DNL_ is about 6 % of the LSB, yielding an RMS value of 2.88 ps: on average the time width of the histogram bins is 48 ± 2.88 ps.

### INL - Integral non-linearity

The INL is used to determine to which degree the response of a time-tagging system differs from the ideal linear behaviour. The INL gives an estimate of the difference between a TTM measurement and the actual time of arrival of a photon. If the TTM is linear, the INL is constant and zero.

The *INL(i*) is computed as the cumulative sum of all the DNL_(*i*)_ contributions, Eq. S4:

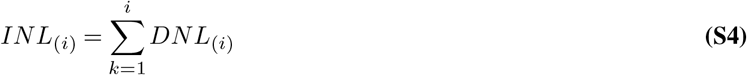

The *σ_INL_* for all the TTM channels is 8 % of the LSB, which corresponds to an RMS value of 3.84 ps.

## Supplementary Note 3: Flash TDC module

In this supplementary section, we describe the principal components and the associated functions that constitute the flash TDC module used in the TTM architecture. The core constituent of a flash TDC module is a tapped delay line (TDL). The TDL is made up by a series of small delay elements joined in a chain architecture and is used to delay an input (START) signal with respect to a reference-sampling FPGA digital clock. From the delay characteristics of each delay element, it is possible to match the distance covered by the START signal along the delay line with a specific arrival time (time-tag) with respect to the FPGA clock signal (8, 9), Fig S8 a.

### Tapped Delay Line and latch barrier

A specific Xilinx FPGA primitive function block known as CARRY (CARR4 or CARRY8 depending on the FPGA family) was used as a fundamental delay block to build the tapped delay line, formed by connecting multiple CARRY blocks. Each delay block is connected to a latch component that can retain either a boolean ’0’ or a ’1’ depending on the data input value at the rising-edge of an FPGA clock. Assuming that the START signal is ’0’ (false) at the steady state and ’1’ (true) after a triggering event, the ’1’ value will propagate through the tapped delay line elements and consequently the data input of each latch (bit_*n*_) will also change from ’0’ to ’1’. The START signal propagates freely until a rising-edge of the FPGA clock occurs. This rising-edge freezes the values of the latch barrier connected to the TDL elements giving out a digital reading of how far the START signal travelled along the TDL within the FPGA internal clock period. Knowing the reciprocal relationship between the CARRY delay value (and also knowing the delay contribution of each CARRY element in the chain) and the distance the START signal covered before the rising-edge of the FPGA clock occurred, it is possible to convert the digital measurement of the travelled distance into a time measurement (10), Suppl. Fig. S9 a.

### Thermometer to binary encoder

The readout from the latch barrier is a series of ’ones’ and ’zeros’ that have to be converted into a binary value for a more efficient representation. Therefore, a dedicated circuit is needed to interpret and decode the TDL data is. The thermometer-like readout coming from the latch barrier is sent to a thermometer to binary encoder component (T2B). The T2B accepts an array of n-bits as input and returns a binary number that represents how many ’ones’ are present in the input latched data. This T2B conversion simplifies the TDL readout, allowing for a more effective data registration in terms of memory resources utilisation (11), Suppl. Fig. S9 b.

---

**Fig. S1.**
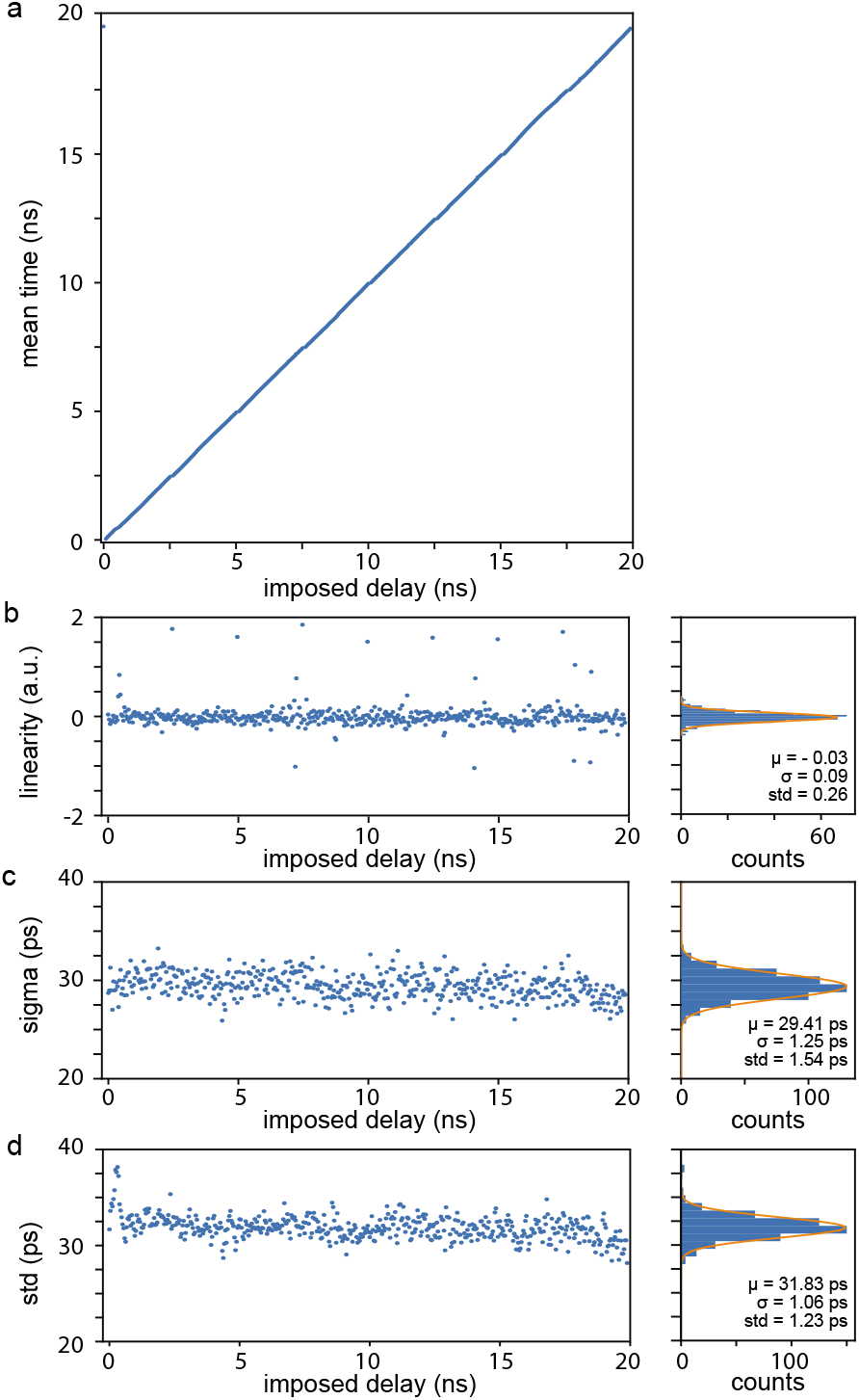
Single Shot Precision Measurement. Detailed representation of the single-shot precision experiment for the photon channel #11. The experiment uses the BrightEyes TTM to measure repeatedly the start-stop interval (i.e., the delay) between a fixed 50 MHz signal, used as SYNC signal, and a synchronised second signal, used as photon signal. The experiment is repeated for all possible delay values (imposed delay) withing the repetition rate of the SYNC signal, i.e., 20 ns. For each imposed delay values, the TTM collects several millions of sync-photon pairs, and builds up the start-stop time histograms (not shown). Each single histogram is fitted with a normal (Gaussian) distribution *A* exp { – ((*t* – *μ*)/*σ*)^2^/2} to extract the mean *μ* and the standard deviation *σ* values. (a) The mean value *μ* as a function of the imposed delay, (b) The difference between the imposed delay and the mean value obtained as function of the imposed delay, (c) The standard deviation *σ* value as function of the imposed delay. (d) Calculated standard deviation of the start-stop time histogram as a function of the imposed delay. The similar values between the Gaussian standard deviation *σ* and the calculated standard deviation demonstrates the normal distribution of the start-stop time histogram. One the right-side of each graph the relative statistics is reported.

**Fig. S2.**
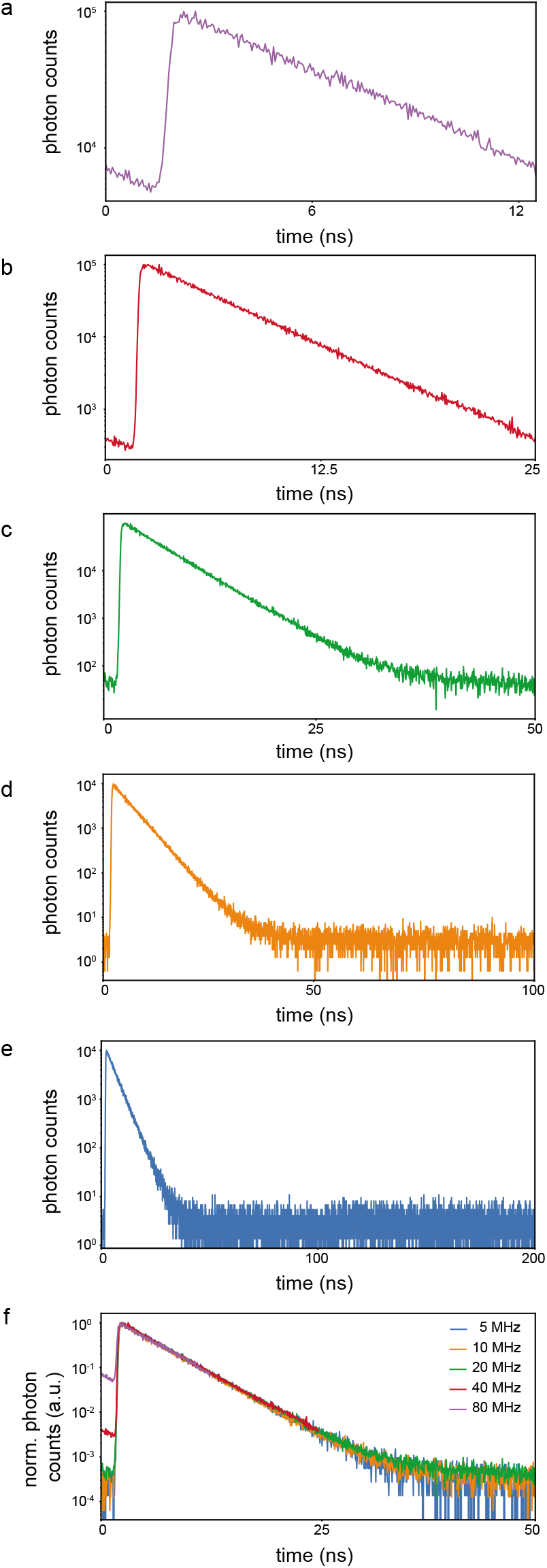
Validation of TTM for different temporal ranges (a-e) and comparison of the obtained results (f). Fluorescence decay histogram, photon counts as a function of time, of a fluorescein-water solution for **a** 80 MHz, **b** 40 MHz, **c** 20 MHz, **d** 10 MHz and **e** 5 MHz laser repetition rates. **f** cumulative view, normalized photon counts versus time, of the reconstructed fluorescein decay histograms for all the probed temporal ranges. 40 MHz and 80 MHz curves show an higher offset when compared with 5, 10, and 20 MHz plots, due to a not complete relaxation of the fluorescein molecules from the excited state that occurs with shorter laser excitation periods.

**Fig. S3.**
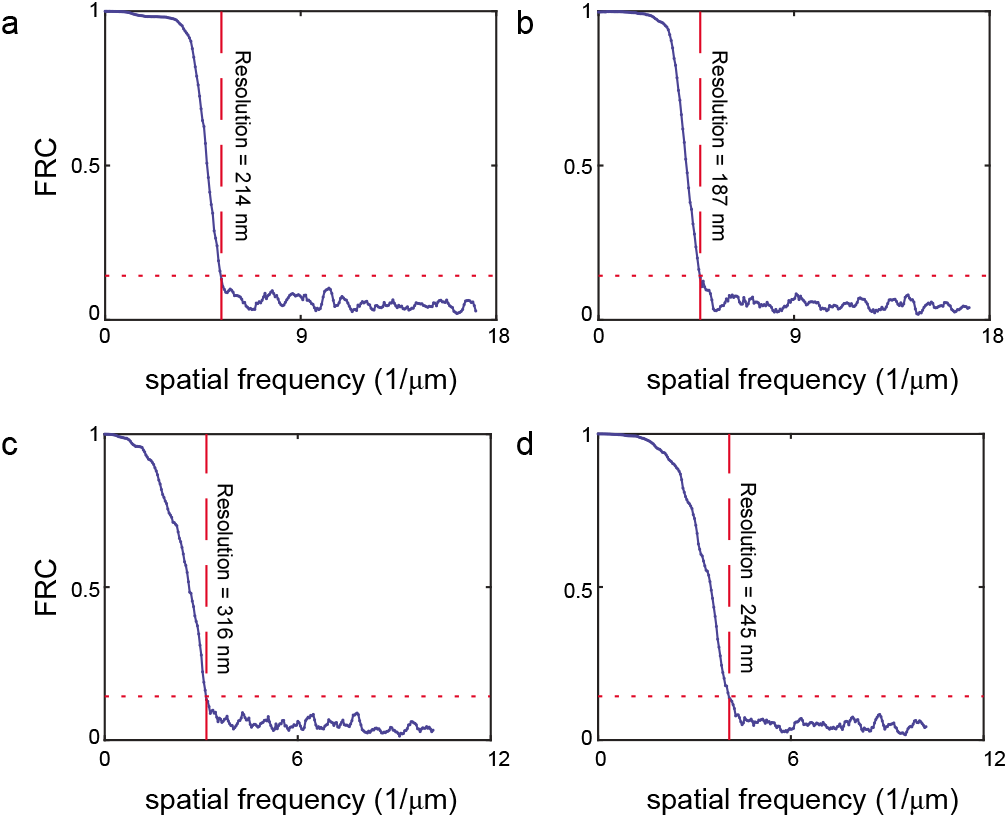
Fourier ring correlation (FRC) analysis for ISM and confocal images of YG carboxylate fluoSpheres (a-b) and fixed-cells labelled for vimentin visualisation (c-d). The analysis compares side-by-side the FRC analysis for the confocal closed pinhole (0.2 AU) images (left) and the APR-ISM images (right). Each curve represents the decay of the correlation as a function of the spatial frequency. We used the 1*/*7 criteria (short dotted red lines) to estimate the effective cut-off frequencies of the images. The inversion of these values, which are reported in each graph, represents the effective resolution of the images.

**Fig. S4.**
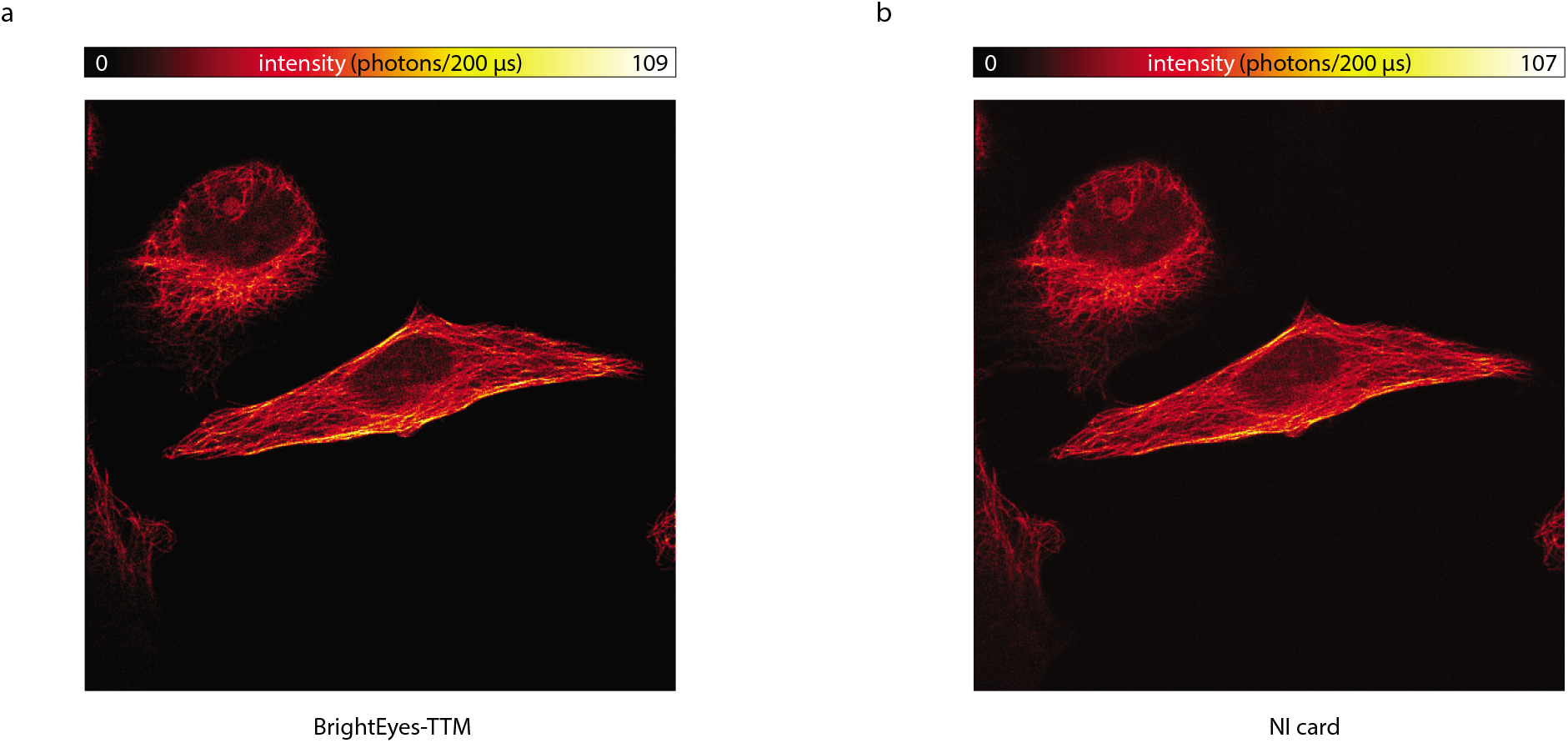
Imaging with the off-line BrightEyes TTM and the real-time NI-DAQ systems. Side-by-side comparison of confocal (0.2 AU) images obtained with our TTM (a) and the NI-DAQ systems (b). The images represent a *α*-tubulin immunolabelled Hela cell. The images are collected simultaneously: The BrightEyes TTM receive the singnal from the central element of the SPAD array detector, duplicated it, sent one copy to the NI-DAQ system, and use the original signal to create the time-tagged data-set. The NI-DAQ system generates in real-time the confocal images, whilst the TTM image is generated off-line.

**Fig. S5.**
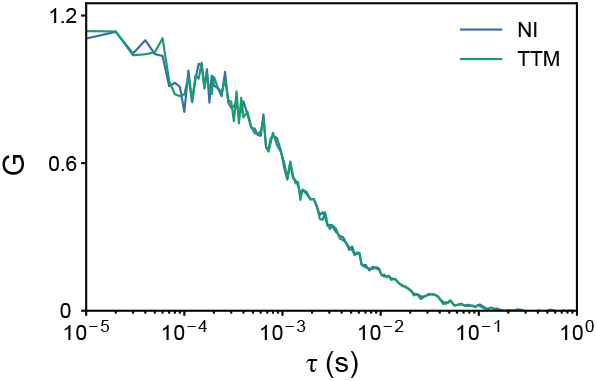
Single-spot fluorescence correlation spectroscopy with the BrightEyes-TTM and NI-DAQ system. Comparison of the two autocorrelation curves obtained recording the simultaneously the signal with the two different platforms. The autocorrelation curves represent the signal collected from the central element of the SPAD array detector, and describe the freely diffusing fluorescent beads. Average over 13 traces of 10 s each.

**Fig. S6.**
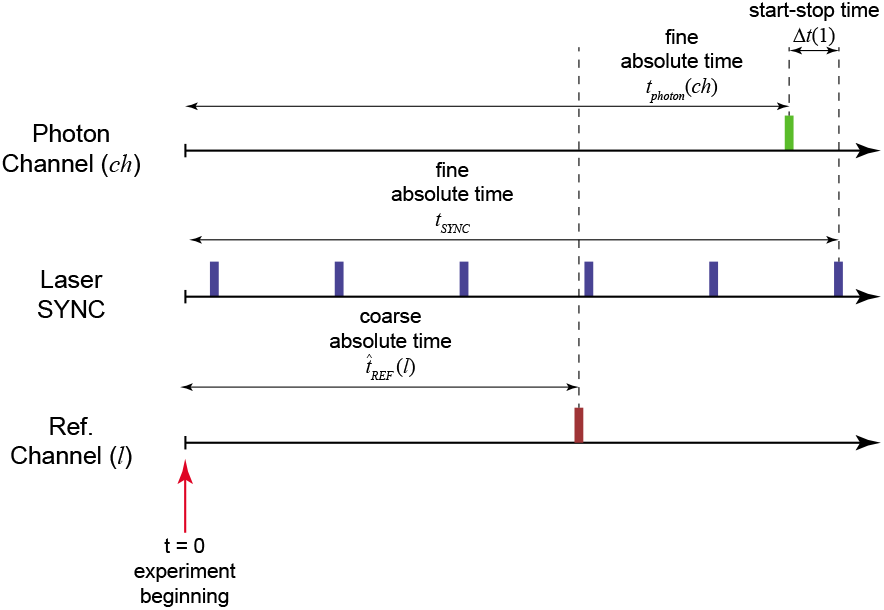
Time-Tagging Principle. The time-tagging (TT) mode allows recording individual events and labelling each of them with a temporal signature. Typically this temporal signature denotes the delay time of the events in respect to the beginning of the experiment (absolute time). Our TT module is able to tag three different class of events: The photon events, i.e., a photon is registered by the detector which delivery a digital signal to the module; The sync laser events, i.e., the synchronisation signal delivered by a pulsed laser; the reference events, i.e., a signal generated by another component of the experimental setup (e.g., an actuator, a laser modulator). Each class of events report the temporal signature with a different precision: in our TTM the absolute time for the photon events (*t*_photon_) and the laser synchronisation events (*t*_SYNC_) have few tens of picosecond precision, and the absolute time for the reference events 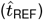 has few nanoseconds precision. Starting from these temporal signatures it is possible to derive many other different temporal information. For example, for each photons event it is possible to derive the so-called start-stop time (Δ*t*), which describe the delay of the photon event in respect to the successive sync laser event. Table 3 describes the main temporal signatures used in this work. Very important, together with the temporal signatures, the TTM records also the number of the channels (*ch*) or inputs (*l*) associated to the photon or reference event. In our work, the channel for the photon event describes the element of the SPAD array detector which collected the photon, thus it can be considered a spatial signature.

**Fig. S7.**
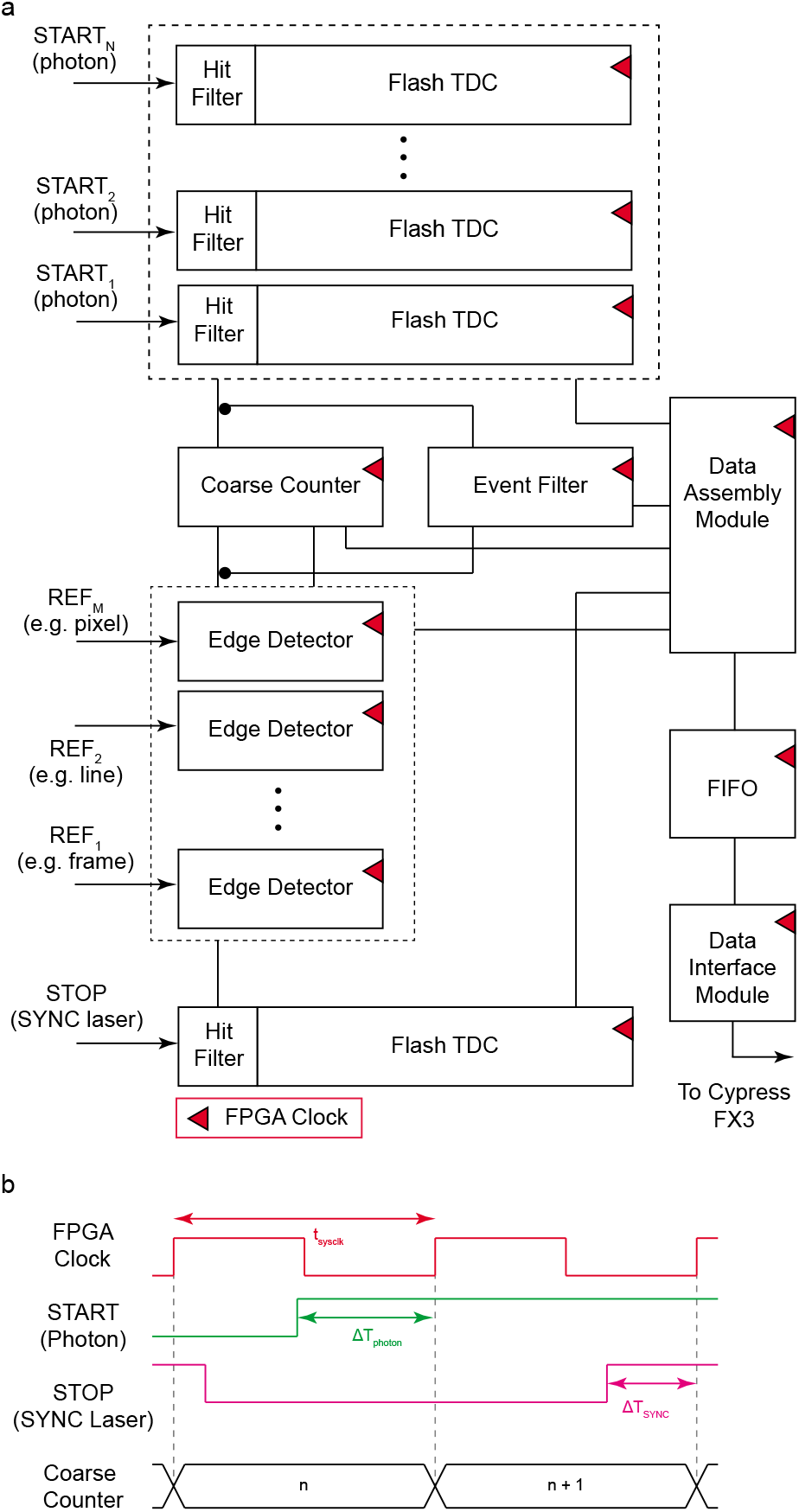
Interpolating TTM-TDC FPGA architecture (a) and general fundamental concept of the sliding scale approach (b). **a** General and simplified circuit architecture of the FPGA-based BrightEyes-TTM. Tapped delay line-based flash TDC modules and relative hit-filters for sampling the START_N_ (N = 21) signals, with picosecond precision, with respect to the FPGA clock (FPGA clock in red) (top portion of the figure). Free running coarse counter for implementing the sliding scale TDC technique and for sampling the REF_M_ (M = 3) synchronization signals with a nanosecond precision (~4,2 ns). Event-filter circuit to reduce the data throughput by transmitting information only when photons are detected (middle-top portion). Edge detector components for sampling external reference signals with nanosecond precision (i.e. pixel, line, frame clock of an imaging SP-LSM system) using a counter-based coarser TDC approach (middle-bottom portion). Single tapped delay line (hit-filter & flash TDC module) for acquiring (with picoseconds precision) the STOP signal shared by all the STARTN inputs (bottom). Data assembly module for collecting a multiplicity of input digital signals and values (i.e. Δ*T*_START_(*ch*) and *n*_photon_(*ch*) as well as valid arrival flags for the photons, Δ*T*_STOP_ and *n*_SYNC_ as well as its valid digital flag for the laser sync, and the valid arrival flags for the REFM signals) and arranging the digital data tags into a suitable form in order to be stored into a FIFO memory, FIFO memory to buffer incoming data awaiting to be sent over a host-processing unit via the data interface module through the Cypress FX3 chip (right). **b** TTM interpolating architecture working principle. FPGA clock used to (i) drive all the TTM architecture components and (ii) as a fundamental reference signal for all the time measurements (top). Representation of how the TTM circuit architecture tags the START-photon (green) and STOP-sync (pink) signals with respect to the FPGA sampling clock (middle) and saves/computes both the Δ*T*_START_ and Δ*T*_SYNC_ (i.e. integer values representing the number of tapped-delays - in the delay line - that the signals have travelled through before the arrival of the FPGA clock rising-edge) signals respectively. Coarse free-running counter increasing its value at each FPGA clock rising-edge event for reconstructing, in a post-processing phase, Δ*t* photon start-stop time.

**Fig. S8.**
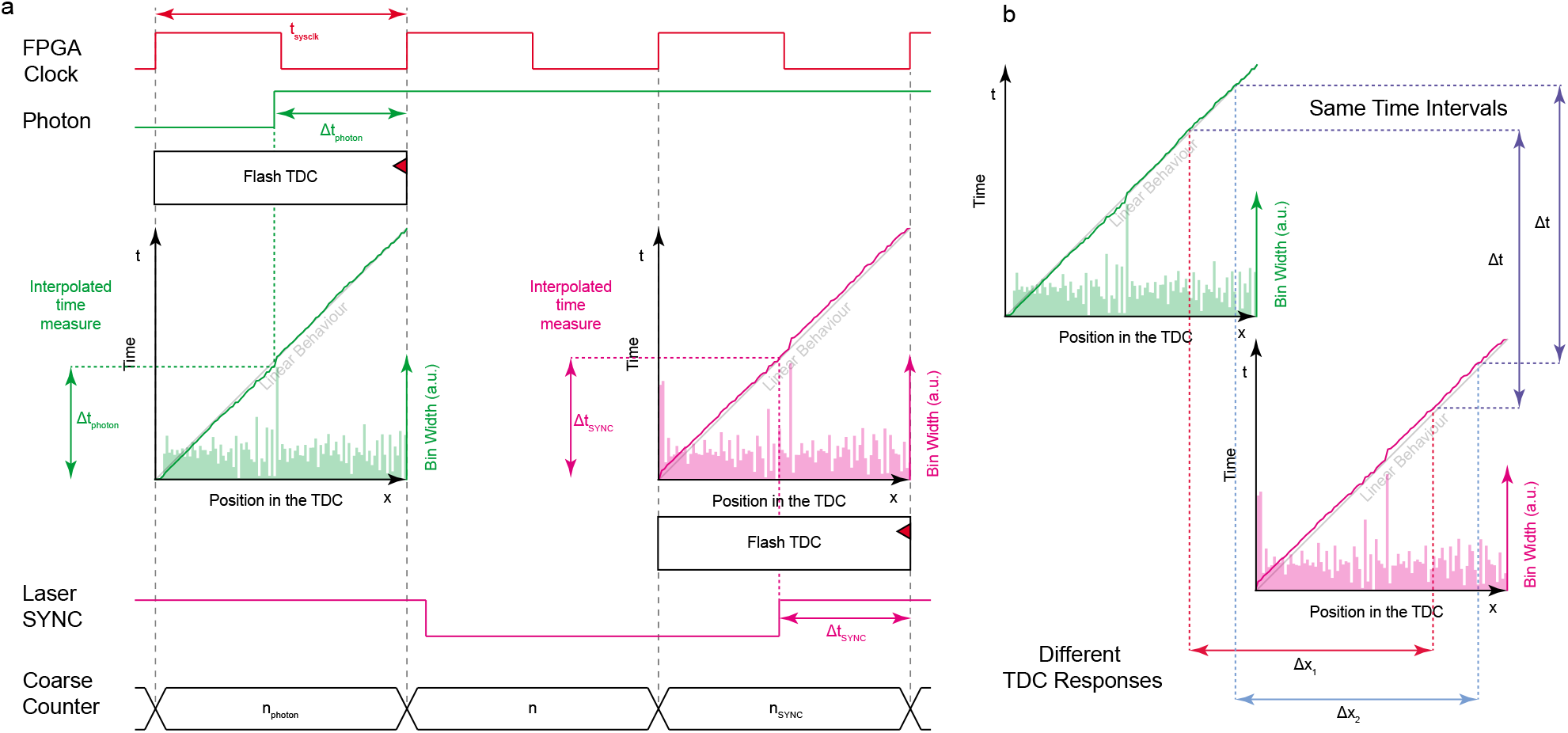
**Schematic of TTM sliding scale technique working principle (a) and mapping of** Δ*t*(*ch*) **start-stop time accross START_N_ and STOP_SYNC_ tapped-delay-lines.** **a** START_N_ (photon) arrival time (Δ*t*_START_(*ch*)) is computed with respect to the rising-edge of the FPGA clock (top) thanks to a dedicated flash-TDC module of which the time response and the tapped-delay-line bin widths are shown as a function of the bin number (left-middle). As soon as START_N_ is detected also the value *n*photon(*ch*) of the coarse counter gets registered (bottom). The STOP_SYNC_ arrival time (Δ*t*_SYNC_) is also recorded on its the dedicated flash-TDC module (time response and bin-widths are show in the right-bottom portion) with respect to the FPGA clock together with the corresponding value of the coarse counter *n*_SYNC_ (bottom). Δ*t*_START_(*ch*), *n*_photon_(*ch*) and Δ*t*_SYNC_, *n*_SYNC_ are used to compute Δ*t*(*ch*) start-stop time according to Suppl. Eq. S1. **b** Smoothing out flash-TDC non-linearity: because of the START_N_ and STOP_SYNC_ signals are asynchronous with respect to the FPGA clock, the same start-stop time interval Δ*t*, (i.e. the delay between a photon and a laser sync event) is measured at different positions in the flash TDC modules.

**Fig. S9.**
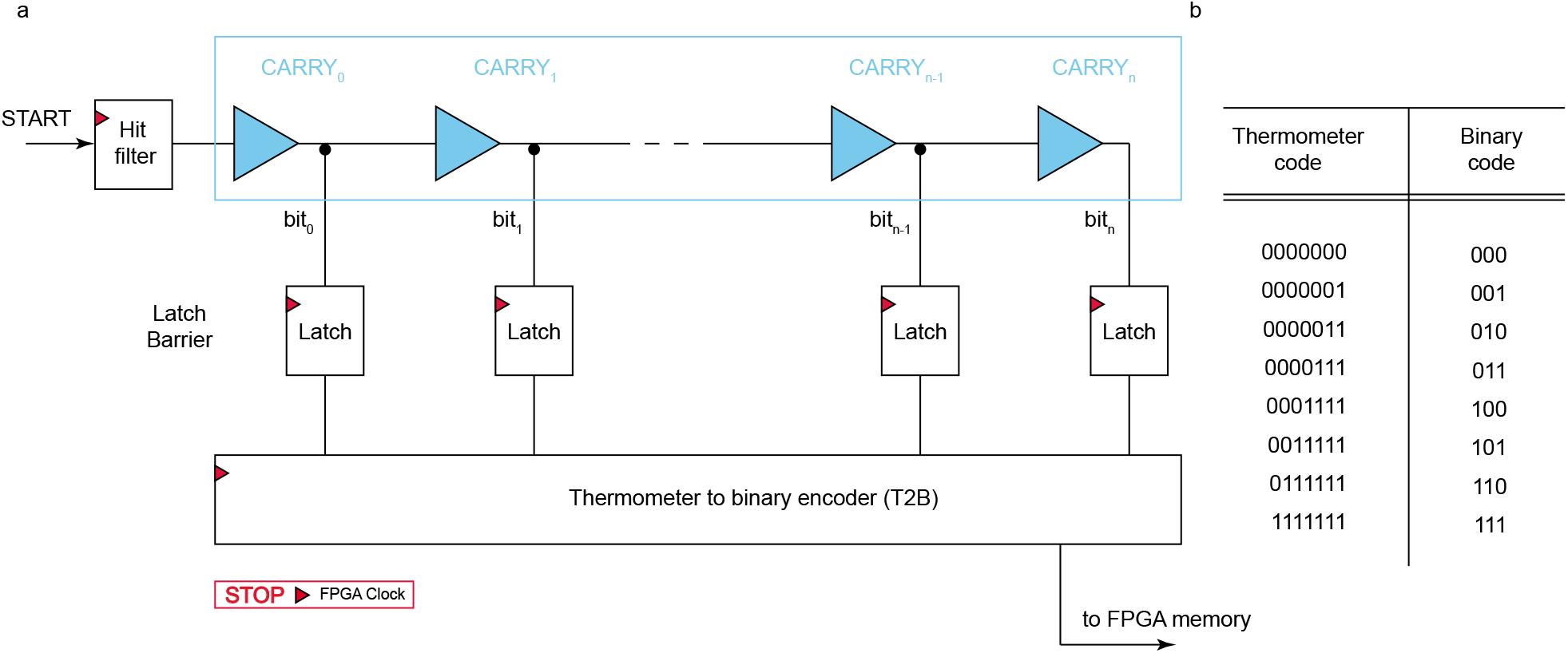
Architecture of the flash time-to-digital (TDC) converter (a) and functionment of the thermomether-to-binary encoder (b). **a** Flash TDC module comprenshive of a hit-filter (Suppl. Fig.S10) and tapped-delay-line made up by inbuilt FPGA CARRY delay elements (top). Latch barrier to sample and stabilize the tapped-delay-line readings (i.e. Δ*T*_START_(*ch*) or Δ*T*_STOP_) (middle). Thermometer to binary encoder to translate the tapped-delay-line readings into a binary form (bottom). **b** Example of thermometer-to-binary working principle: tapped-delay-line readings are converted into binary number for a more compact and efficient data representation.

**Fig. S10.**
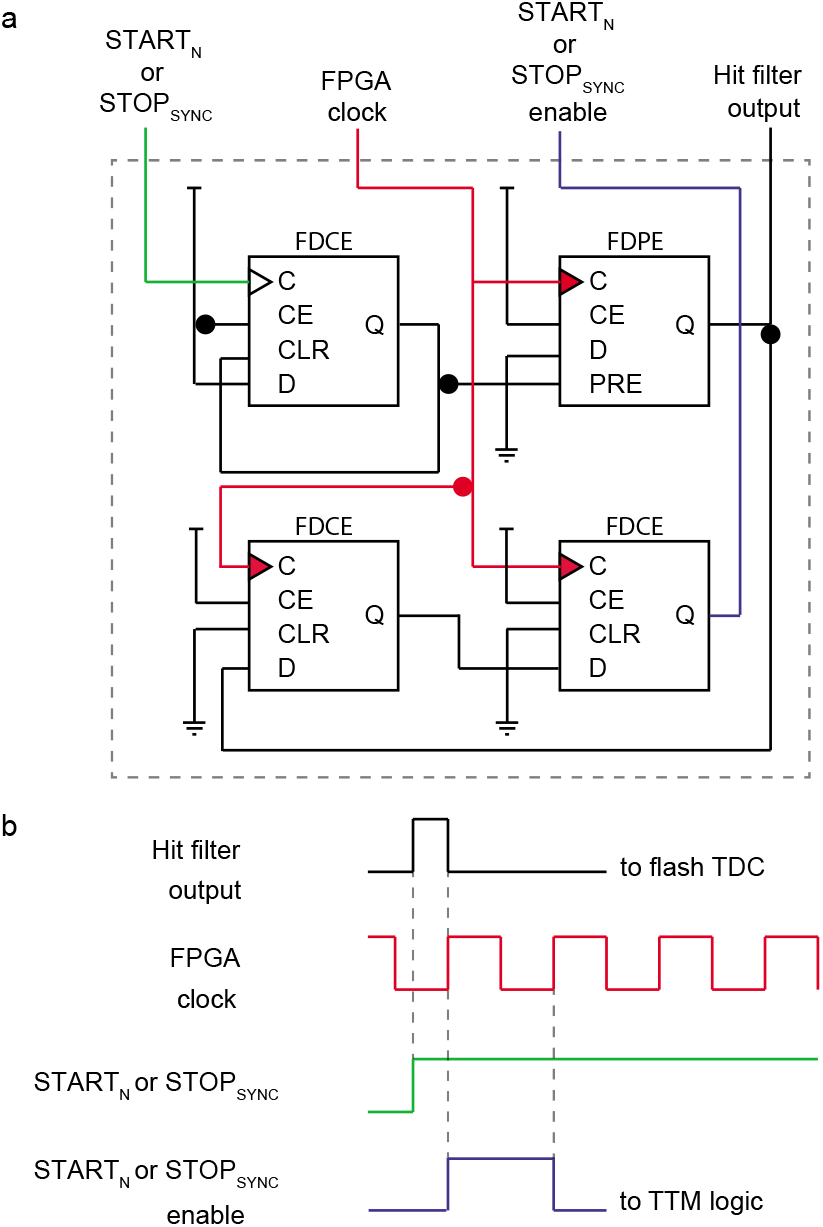
Circuit schematic of the hit filter component (a) and related input-output digital signals (b). **a** Hit-filter FPGA digital circuit. The intimate digital-electronic layout of the hit-filter made up by only four flip-flops. The hit-filter component is engineered and deployed to shape the incoming photons and sync signal lengths based on the sampling FPGA clock period and, at the same time, for generating a toggle signal event (the photon or sync enable signal) for each detected STARTn (photons) and STOP (sync) events. The hit-filter logic is also necessary to avoid the clogging of the flash TDC module and allow for the TDC module to be ready to sample incoming signals thus reducing the dead-time of the architecture which, thanks to the hit-filter, becomes independent from the signal pulse duration (hold-off). While the hit filter output signal is solely used to activate the flash TDC module the photon enable signal is distributed to the entire TTM logic to sample and record the arrival of a photon (or sync pulse) on a specific channel. **b** Input and outputs signals of the hit-filter logic. Primary digital hit-filter output (top), sampling FPGA clock (second row), rinsing edge of photon or sync event (third row), photon or sync enable signal that has the duration of an FPGA clock period (bottom).

**Fig. S11.**
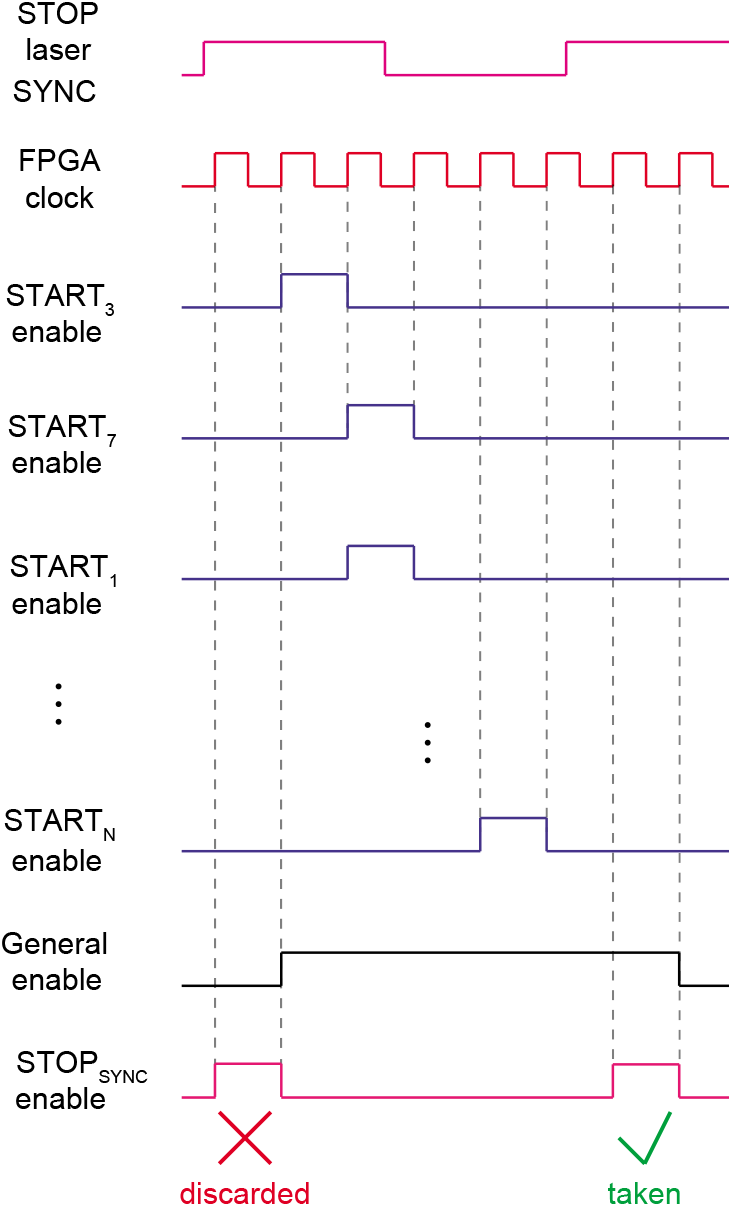
Circuit logic of the event filter component. All photon valid-enables signals (from START_1_ enable to START_N_ enable), produced by the hit-filter modules whenever a START-photon (or STOP_SYNC_) event is detected, are used to efficiently record the multi-channel data tags (Δ*T*_START_(*ch*) and Δ*T*_STOP_). The event filter was designed and implemented to optimally handle time-tag data for high repetition rates of the laser sources (up to 80 MHz - Suppl. Fig. S2) used in SP-LSM applications. It is not efficient to sample and tag all the incoming laser pulses (laser SYNC events) and stream the related information together with the data tags (Δ*T*_START_(*ch*), Δ*T*_STOP_, *n*_photon_(*ch*), *n*_SYNC_) of sampled photons to a processing unit. So the event filter circuit works backwards: when a photon is detected in channel i, the corresponding START_i_, enable activates a general enable signal. The general enable signal remains active until the detection of a successive laser pulse sync signal (STOP_SYNC_ enable). If the general enable signal is high at the moment of a STOP_SYNC_ pulse, at least one photon has been detected and only in that case the FPGA circuit tags and registers the START-STOP times of all photons in all channels. By discarding STOP_SYNC_ events that do not have a corresponding START_i_ Enable event, the data rate can be significantly reduced.

**Fig. S12.**
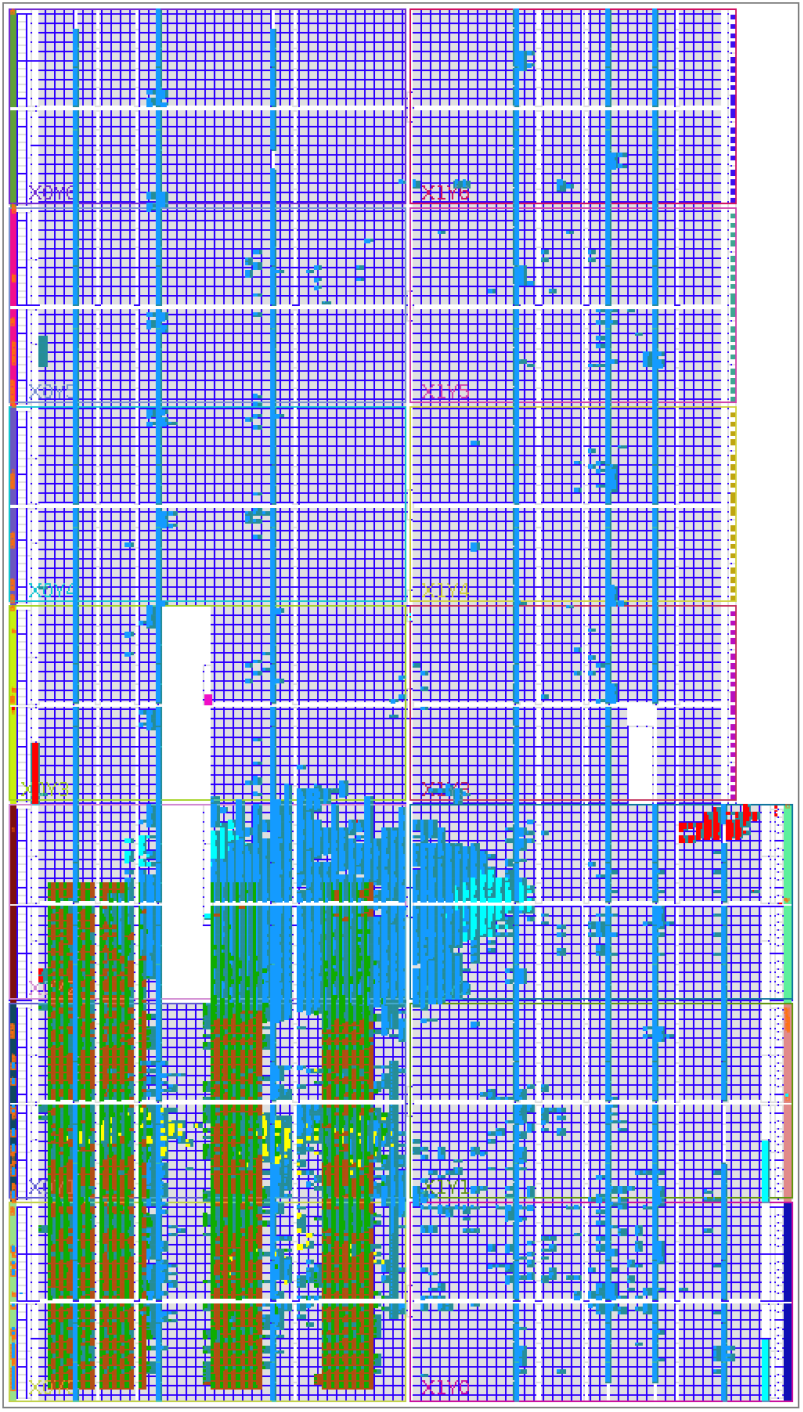
FPGA resource utilisation. 20% of slice look-up-tables (LUT) for Tapped-delay lines (TDLs) (dark red) and thermomemer-to-binary encorders (T2B) in green; FIFO (98 % of inbuilt BRAM) and 5% of LUT for FX3 module in light-blue; Integrated logic analyzer (ILA) debugger in yellow; SYLAP (only for test purpose) in red - 2% LUT.

**Fig. S13.**
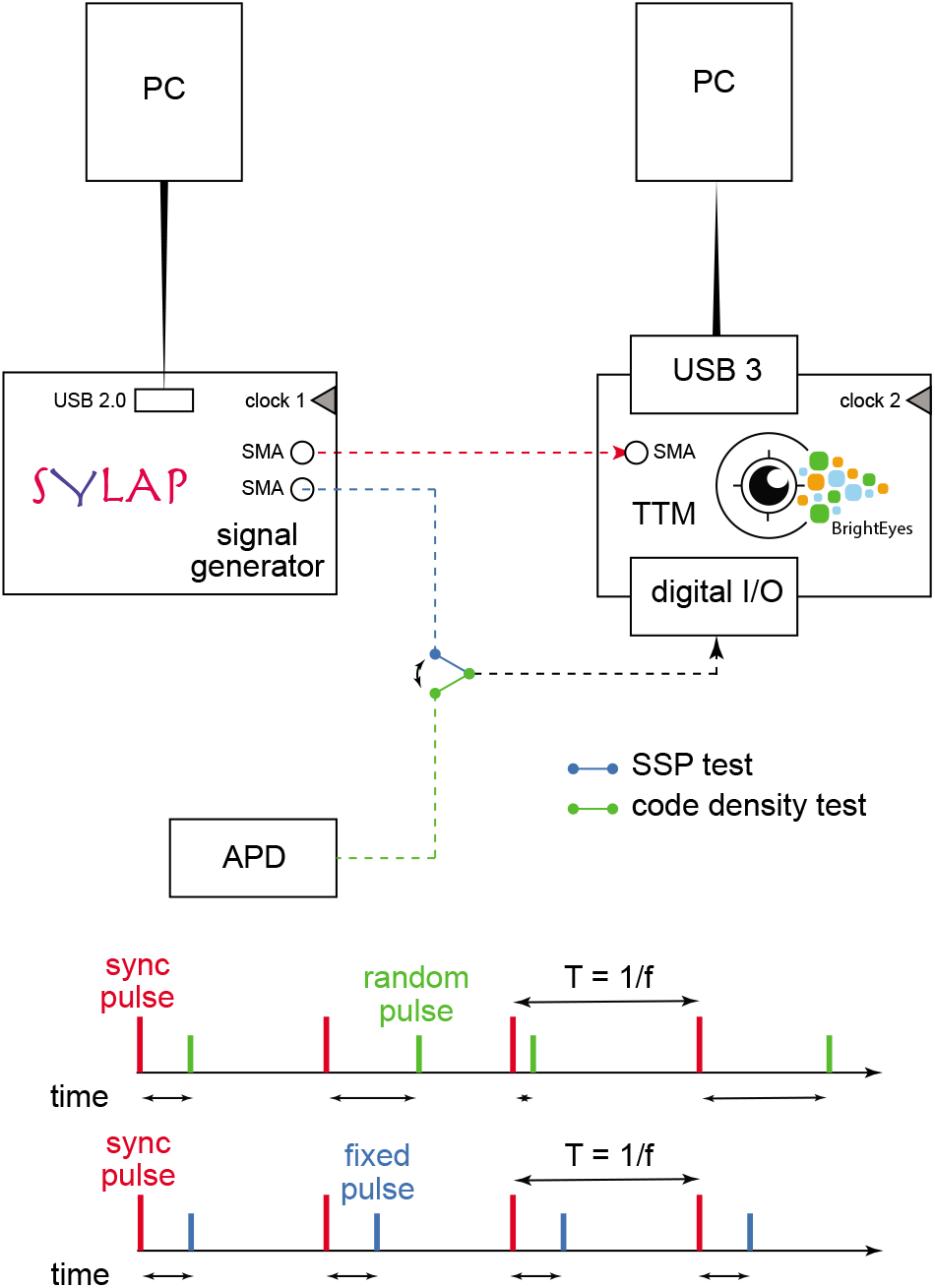
Test Bench System. The setup used for the TTM tests and characterisations. The TTM is STOP connected to clock generated by SYLAP. The TTM start input can be connected the APD for generating asynchronized pulses **(green)**; or can be connected to SYLAP to generate pulses with a fixed delay respect to the clock **(blue)**.

**Fig. S14.**
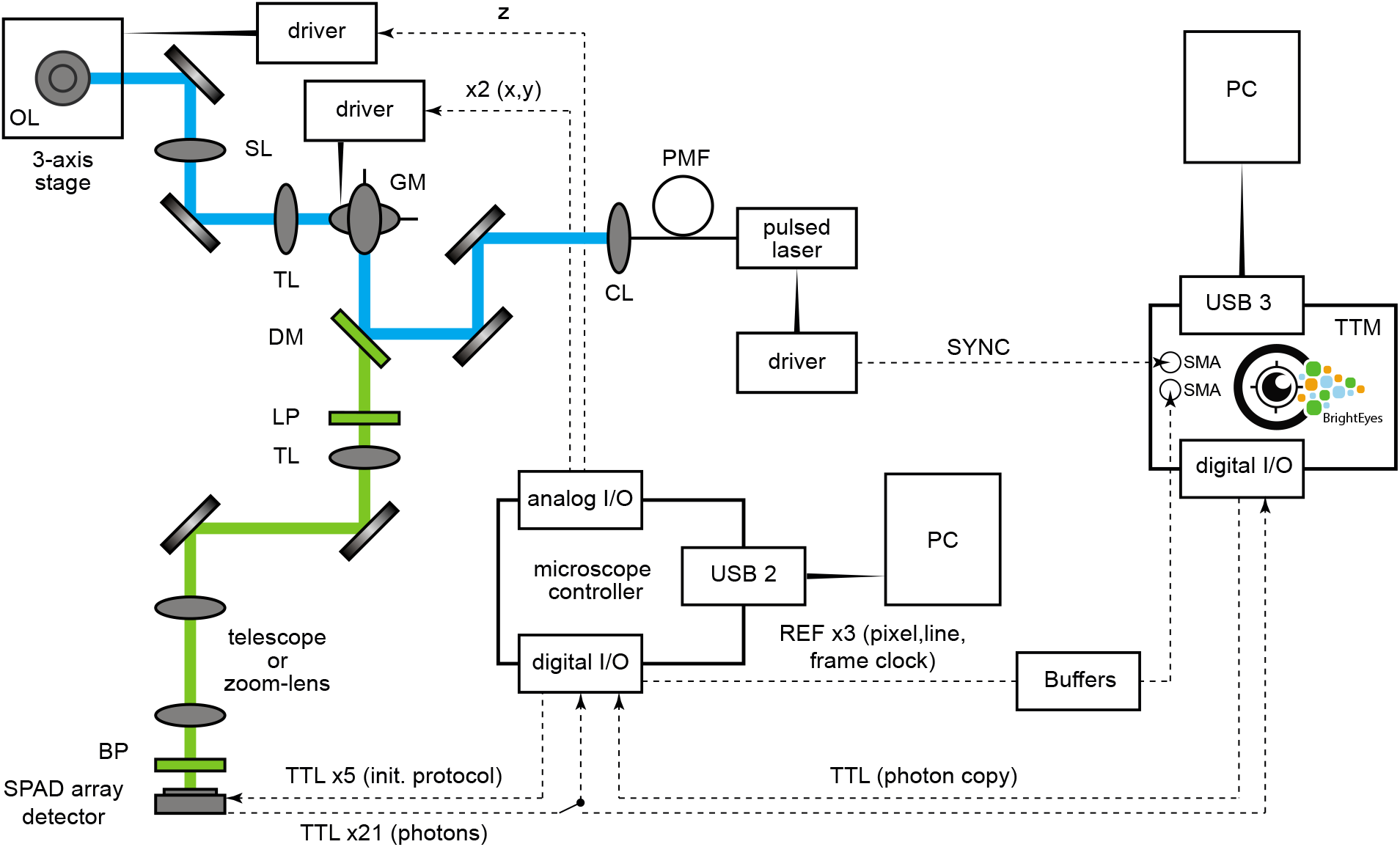
Single-photon laser scanning microscope. Schematic representation of the optical architecture and the data-acquisition and control system. Digital and analogue single-cable connections are represented by dashed lines. DM: dichroic mirror, GM: galvanometric scan mirrors, SL: scan lens, TL:tube lens, OL: objective lens, LP: long pass filter, BP: band pass filter, CL: collimating lens, PMF: polarising maintaining fibre. The pixel, line, and frame reference signals and the laser sync signal are plugged directly to the FPGA-development board using the SMA user I/Os, whilst the photon signals are plugged to the board by using the I/Os daughter card. The board duplicates the photon signal from the central element of the SPAD array detector and send to the microscope controller via a TTL digital signal.

**Fig. S15.**
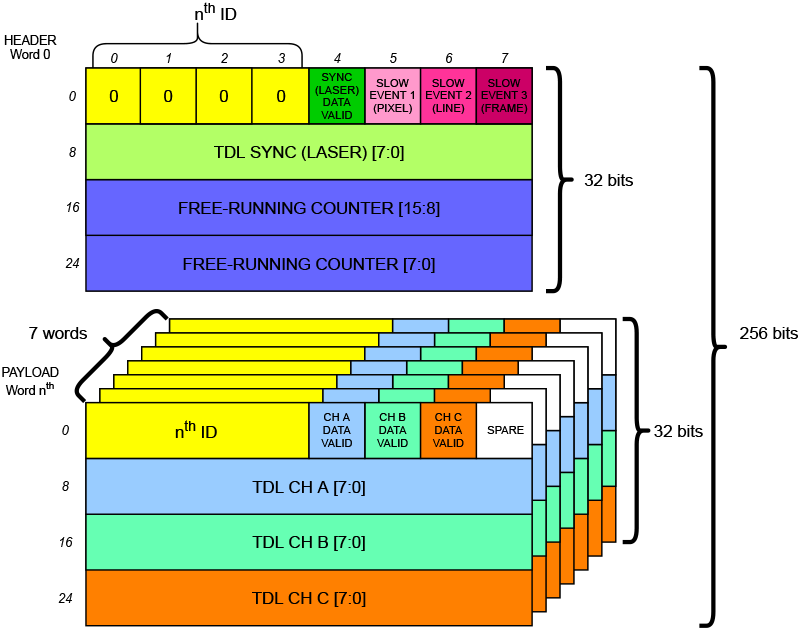
Data Structure. The data structure has an header 32-bit word and, in case of 21 channels, seven payload 32-bits words each one identified with an associated ID number. The channel CH A, B, C are different in each payload words and they are sequentially corresponding to channels 1 … 21.

**Fig. S16.**
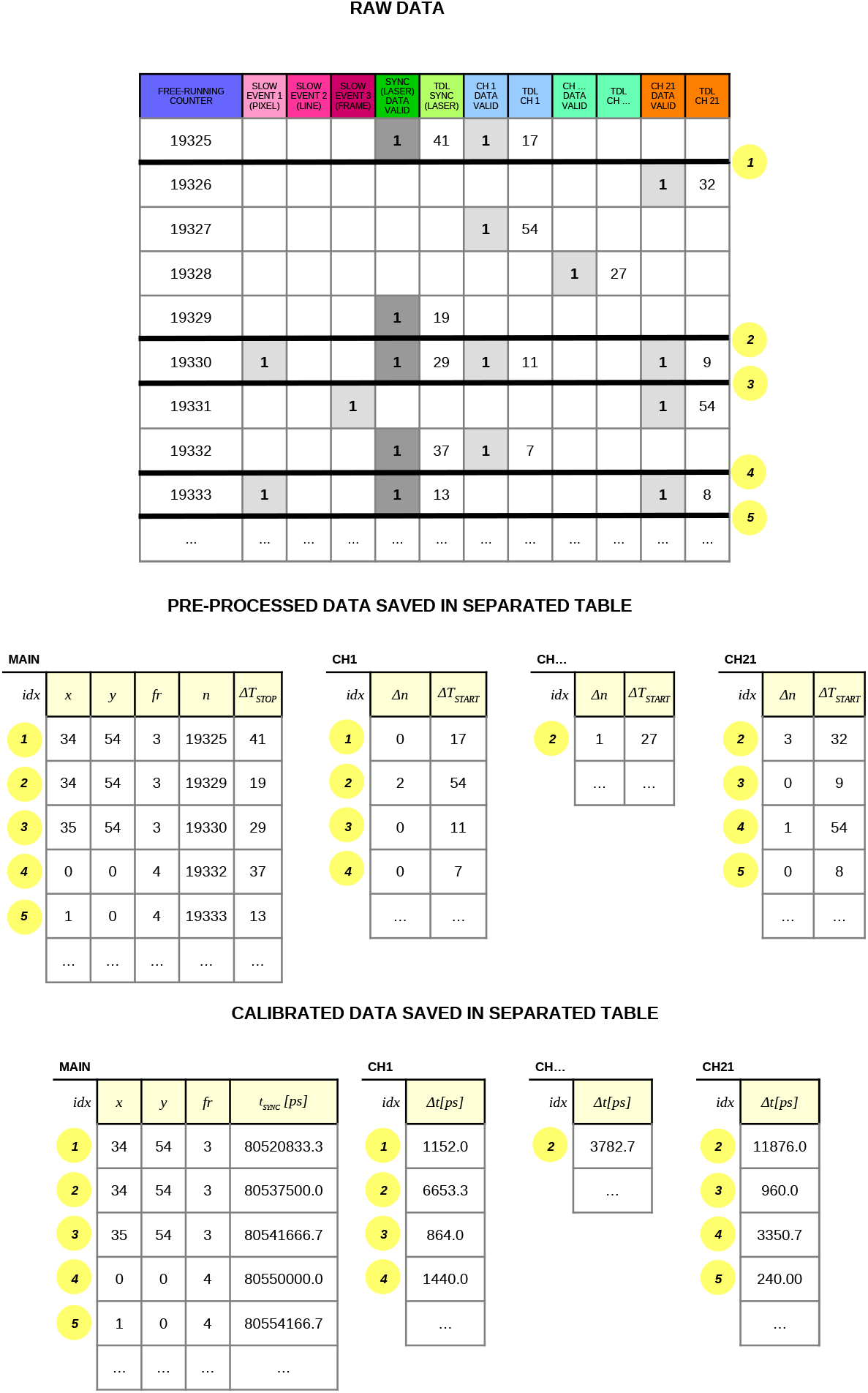
Data pre-processing. The top table represents the raw data as received. In the top table, the gray cells are the data-valid flags. Each SYNC data-valid flags (in dark gray) is associated to a new unique index (number yellow highlighted). The middle tables (main and channels tables) represent the data after the pre-processing. Notably, the channels tables have not all indexes, as they contains only valid events. The lower tables the data are calibrated.

**Fig. S17.**
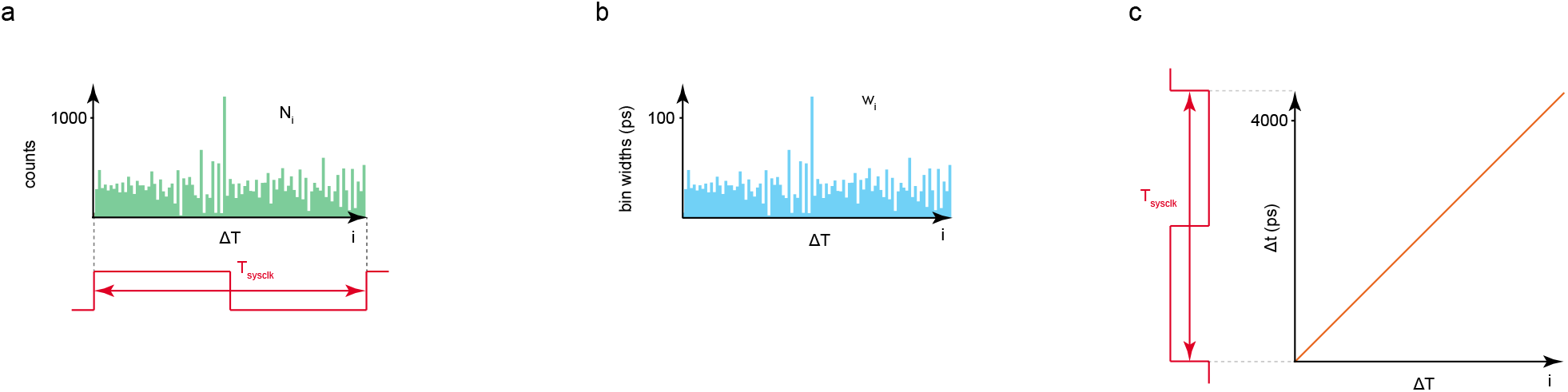
Fundamental steps of the bin-by-bin calibration procedure. **a** tapped-delay-line histogram of received counts as a function of the arrival tap (delay) number. The range of the maximum arrival tap number depends on the reference FPGA clock period 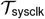: the tapped-delay-line is dimensioned to have a total delay value of 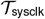. **b** individual value w_i_ of each delay element is estimated (bin time-widths). **c** time as a function of Δ*T*: each w_i_ is used to compute the Δ*t*_START_(*ch*) or Δ*t*_SYNC_

**Fig. S18.**
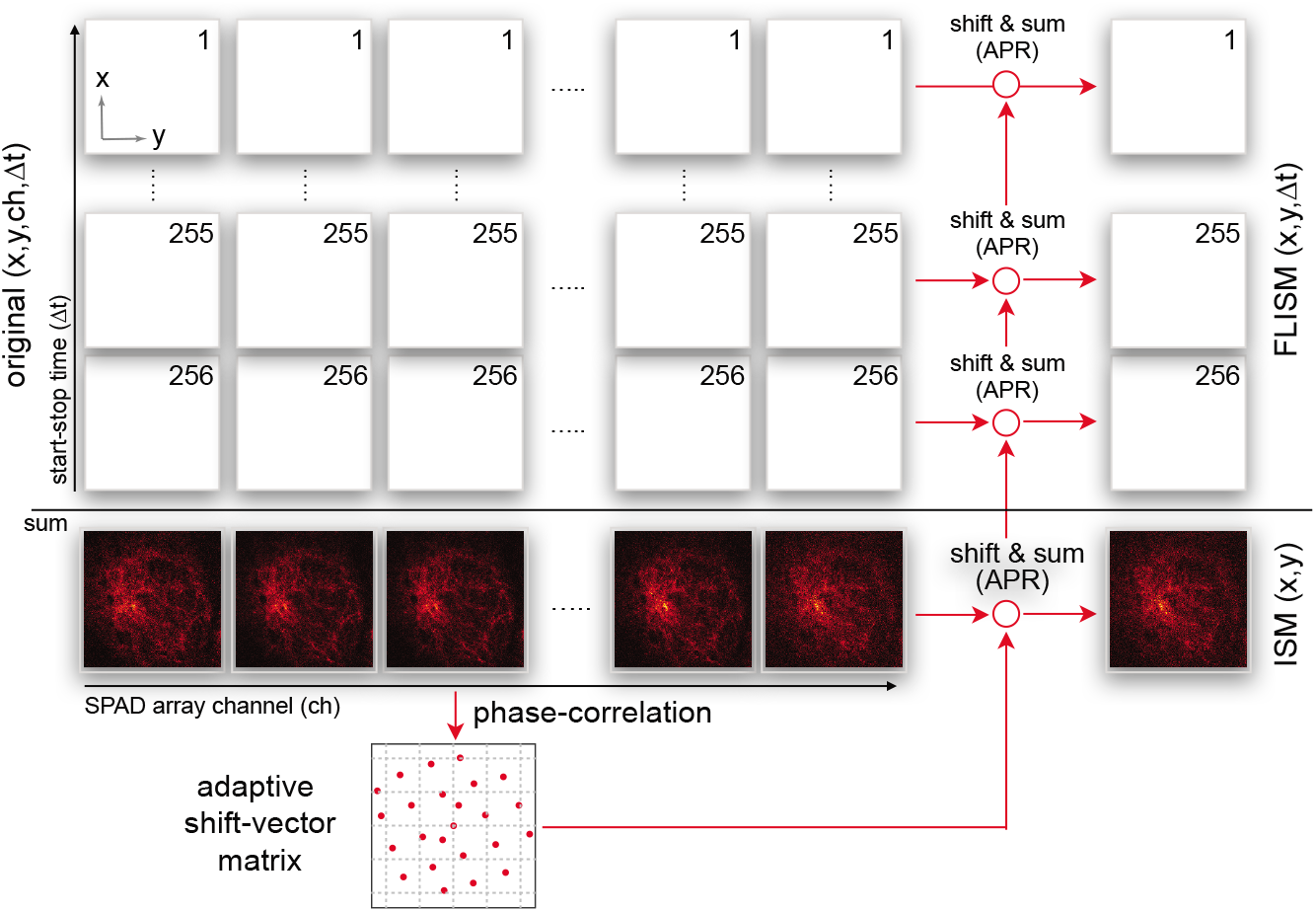
Adaptive pixel-reassignment method for 4D lifetime data-set. In case of imaging, the BrightEyes-TTM provides a 4D (*ch, x,y,τ*) pre-pocessed data-set, which can be used to reconstruct the high-resolution and high signal-to-noise ratio FLISM image. The reconstruction consists in the following steps. (i) The algorithm integrates the data-set along the start-stop *τ* dimension, and generates a 3D (*ch,x,y*) data-set; (ii) a phase-correlation registration algorithm uses the conventional intensity data-set to calculated the shift-vector fingerprint (*s_x_*(*ch*), *s_y_* (*ch*)); (iii) for each *τ* the images (*x, y*) associated to each channel *ch* are shifted according to the shift-vector fingerprint ans summed all together (i.e, along the *ch* dimension). The result is a 3D (*x, y,τ*) which can be used to calculate the lifetime map or the phasor-plot histogram.

**Fig. S19.**
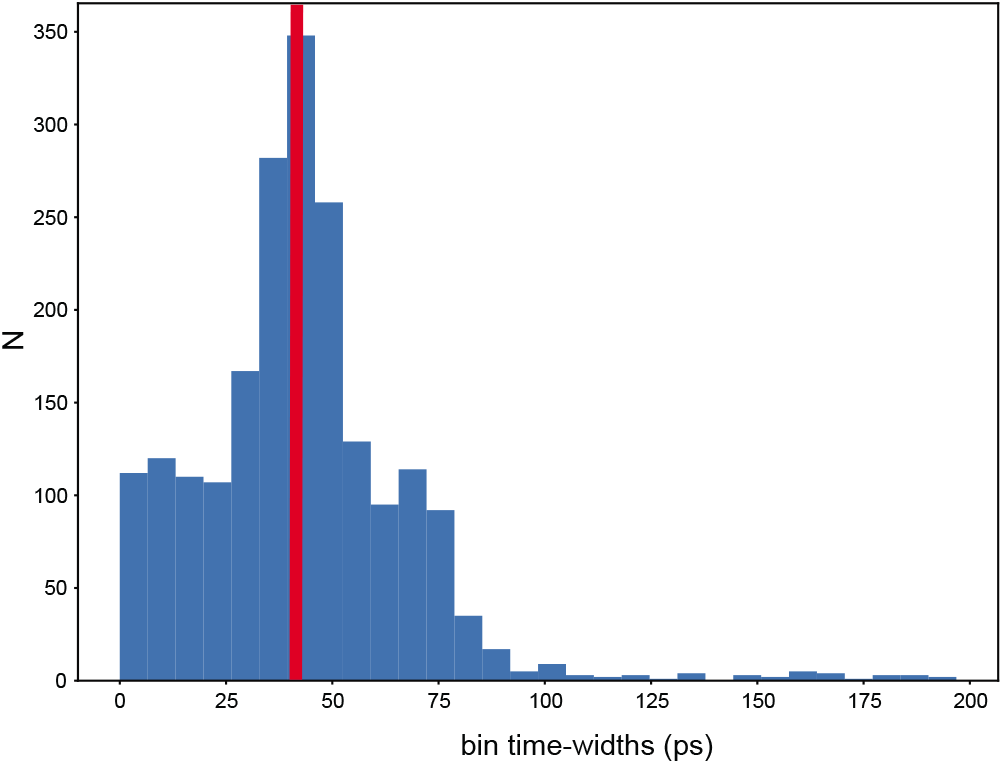
Histogram of the time-bin widths (wi) of all the deployed TDLs in the TTM architecture. Cumulative histogram, number (N) as a function of time-bin widths, of all the calculated w_i_ for all the TTM delay lines. The histogram mean value (red line) is *w*_avg_ = 42.98 ± 25.27 ps.

**Fig. S20.**
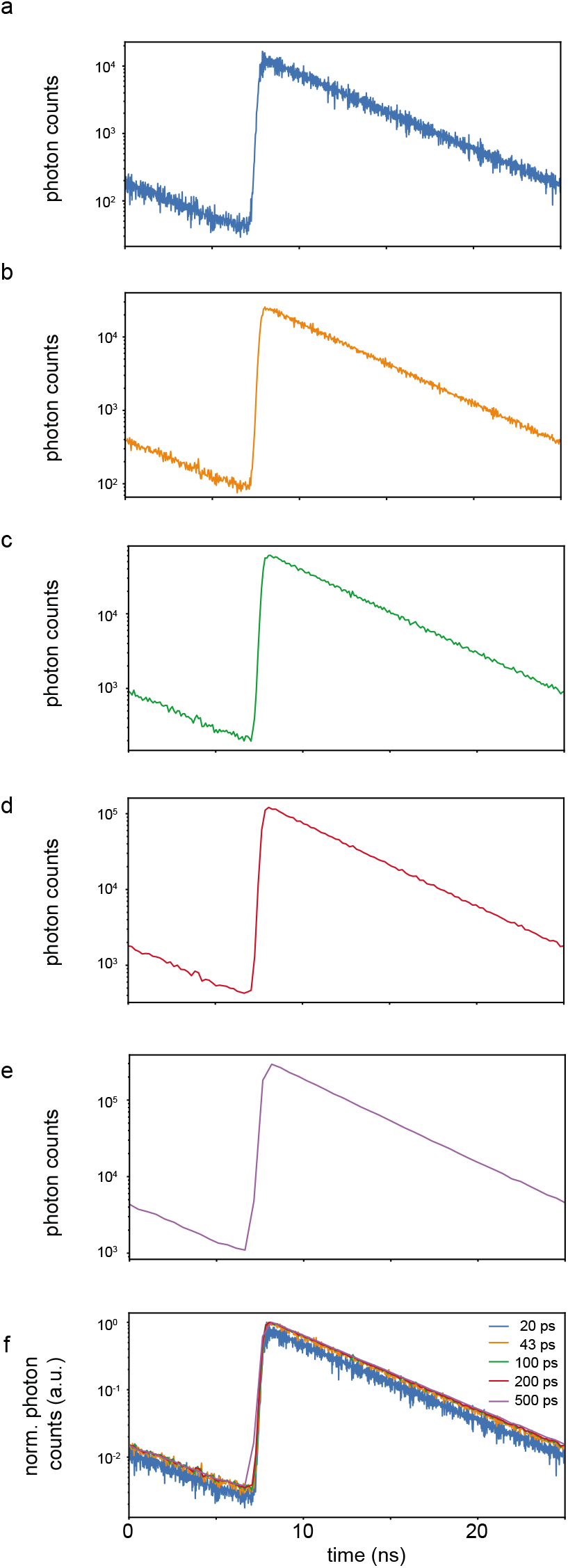
TCSCP histograms of a fluorescein solution reconstructed using different time bin widths (a-e) and superposition of the obtained results (f). Fluorescence decay histogram, photon counts as a function of time, of a fluorescein-water solution reconstructed with bin-width of **a** 20 ps, **b** 43 ps, **c** 100 ps, **d** 200 ps and **e** 500 ps. **f** cumulative view, normalized photon counts versus time, of the reconstructed fluorescein decay histograms for all the different tested time bin-widths. The smaller the time bin-width the higher the SNR, 20 ps curve shows a noisier profile when compared to 43, 100, 200 and 500 ps curves, due to an undersampling of the actual bin-width for the reconstructed TCSPC histograms.

**Table 1.**
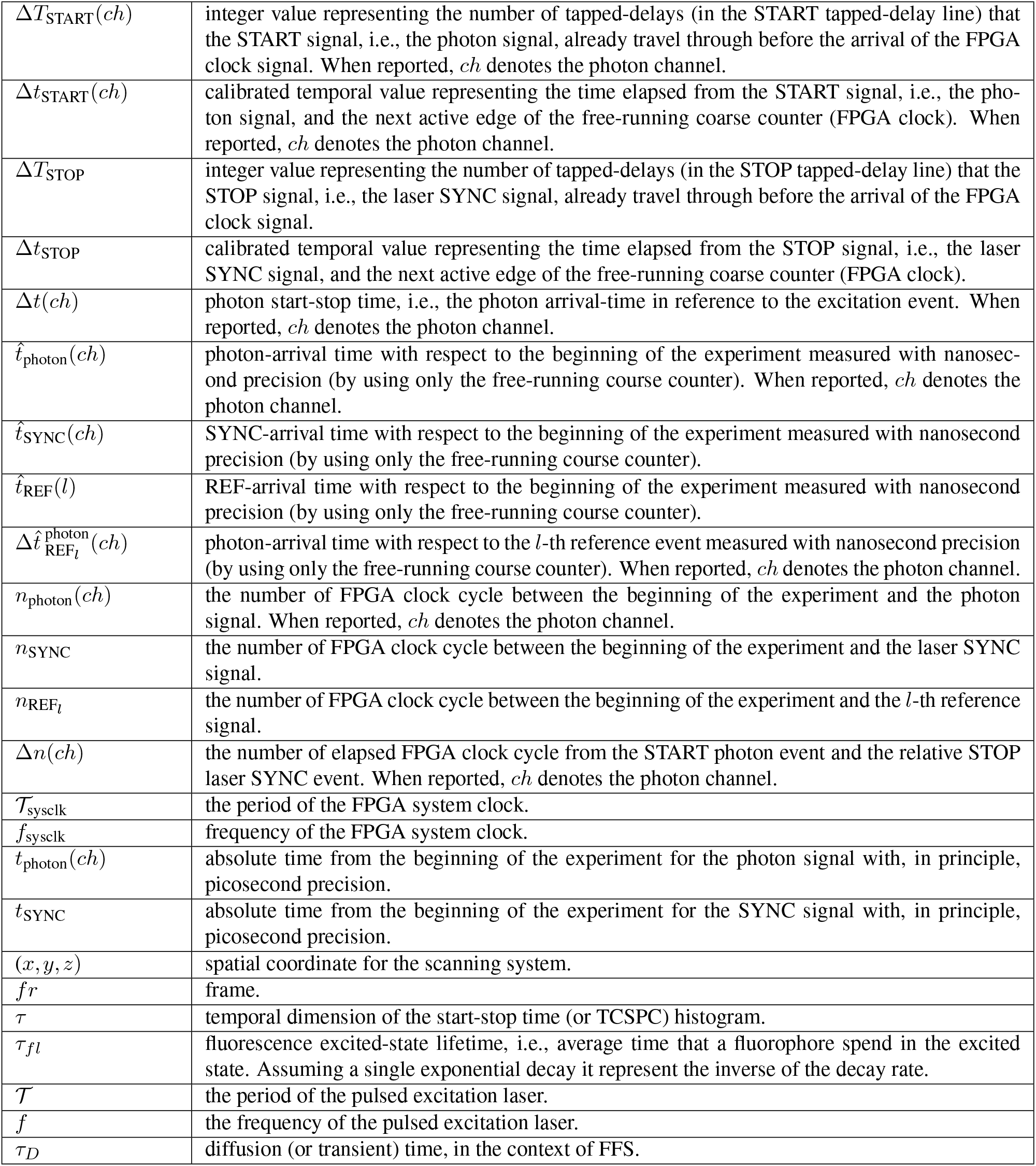
Notations.

